# Speckle-tracking echocardiography in preclinical mouse models: Implications for cancer therapy-related cardiac dysfunction from inhibition of the extracellular signal-regulated kinase 1/2 (ERK1/2) cascade

**DOI:** 10.1101/2025.05.16.654397

**Authors:** Eugene Duvenage, Susanna T.E. Cooper, Daniel Gattari, Joshua J. Cull, Philip R. Dash, Debora Chan, Mariano Llamedo Soria, Mario Rossi, Slawomir J. Nasuto, William Holderbaum, Peter H. Sugden, Peter E. Glennon, Angela Clerk

**Affiliations:** School of Biological Sciences, University of Reading, UK; Institute of Developmental and Regenerative Medicine, Department of Physiology, Anatomy and Genetics, University of Oxford, UK; Functional Genomics and Data Science Laboratory, Instituto de Investigaciones en Medicina Traslacional (IIMT), Universidad Austral-CONICET, Pilar, Buenos Aires, Argentina; Facultad de Ingeniería, Universidad Austral, Pilar, Buenos Aires, Argentina; Grupo de Invesitgación y Desarrollo en Ingeniería Biomédica, Departamento de Electrónica, UTN FRBA, Universidad Austral, Pilar, Buenos Aires, Argentina; University Hospitals Coventry & Warwickshire, Coventry, UK

**Keywords:** Echocardiography, mice, speckle-tracking, angiotensin II, dabrafenib, trametinib

## Abstract

**Aims:** Echocardiography is used widely in preclinical mouse studies, but the emphasis remains the histological/pathological/biochemical changes in the myocardium. Analysis/reporting of cardiac function is generally limited, often relying on M-mode assessment of a single plane across the left ventricle. The aim was to determine if global/segmental endocardial speckle-tracking (strain) has greater potential to assess function by distinguishing between mouse lines and identifying regional effects of different drugs.

**Methods and results:** Echocardiograms from male C57Bl/6J (commercially-available) or C57Bl/6(R) (bred in-house) mice treated with vehicle or angiotensin II (AngII; 0.8 mg/kg/d, 7 d) were analysed. Global strain demonstrated different degrees of hypertrophy induced by AngII in the different lines, and variation in function (e.g. stroke volume/cardiac output were reduced in C57Bl/6J mice only, with no effect on global longitudinal strain). For C57Bl/6J mice, segmental strain (radial/longitudinal peak displacement/velocity/strain/strain rate) and frame-to-frame analysis of radial/longitudinal displacement identified more significant effects of AngII in the basal/mid-regions of the left ventricle. Thus, speckle-tracking distinguished between responses in different mouse lines. The methodology was applied to studies of anti-cancer drugs that inhibit extracellular signal-regulated kinase 1/2 signalling. Dabrafenib and/or trametinib inhibited AngII-induced cardiac hypertrophy, but dabrafenib alone caused abnormalities in endocardial movement whilst, for trametinib with AngII, more abnormalities were detected than with AngII alone. For both, the dominant effect was in the basal/mid-regions of the left ventricle.

**Conclusion:** Echocardiography in preclinical studies can be exploited using endocardial segmental strain for greater insight into how drugs affect cardiac function in different regions of the left ventricle.

**Translational perspective:** Echocardiography is used widely in preclinical mouse studies, but analysis/reporting of cardiac function is generally limited. This study determines the benefits of endocardial speckle-tracking (strain) to assess cardiac function in mice. Global and segmental strain distinguished between mice of different origins in their response to angiotensin II (AngII), and anti-cancer drugs suppressed the increase in AngII-induced LV mass but did not normalise function. The greatest effects were in the basal/mid-regions of the left ventricle. Endocardial strain analysis of mouse echocardiograms provides valuable insight into how drugs affect cardiac function, maximising information from preclinical studies for understanding of the clinical setting.

## 1. Introduction

Hypertension affects hundreds of millions of people and is a major risk factor for heart failure.^1^ Terminally-differentiated contractile cardiomyocytes hypertrophy (i.e. increase in size and contractile apparatus) to accommodate the increase in workload as the heart initially adapts, but vascular dysfunction and accumulating fibrosis exacerbate the stress, eventually leading to heart failure.^2^ Heart failure is often considered as a decline in left ventricle (LV) systolic function and reduced ejection fraction (EF), but the inability of the LV to pump blood to the body may result from diastolic dysfunction and heart failure with preserved ejection fraction. A hypertensive heart exhibits varying degrees of pathology with features of concentric remodelling, concentric left ventricular hypertrophy or eccentric left ventricular hypertrophy.^3^ Concentric remodelling and hypertrophy resulting from hypertension is often associated with preserved EF.^4^ Efforts to identify novel therapeutic targets for hypertensive heart disease are underpinned by preclinical studies in rodents. Many use angiotensin II (AngII) as a physiologically relevant model of hypertension, but select from a range of AngII concentrations. High concentrations (>1.44 mg/kg/d) are associated with increased mortality from blood vessel rupture,^5,6^ whereas moderate concentrations (∼0.8 mg/kg/d) increase blood pressure gradually (i.e. a slow-pressor dose), reducing risk of blood vessel rupture.^7,8^ Notably, cardiac pathology and dysfunction vary according to concentration, with higher doses having a greater effect on LV hypertrophy and cardiac dysfunction.^9^

The global burden of cancer, like hypertension, continues to increase but, for cancer, many new and effective treatments are becoming available.^10^ Almost all recently developed anti-cancer drugs have specific targets, often key intercellular or intracellular signalling proteins such as protein kinases. However, these drugs are rarely specific and may inhibit wild-type enzymes in addition to having off-target effects. For example, the extracellular signal-regulated kinase 1/2 (ERK1/2) cascade (comprising RAF kinases that signal to MEK kinases that activate ERK1/2) promotes cancer.^11^ Mutations in BRAF (most commonly V600E) drive many cancers and inhibitors such as dabrafenib are targeted to BRAF(V600E).^11,12^ However, dabrafenib inhibits wild-type RAF kinases, in addition to some other kinases. Furthermore, low concentrations of some inhibitors targeting BRAF(V600E) paradoxically activate ERK1/2 at low concentrations because they bind to wild-type RAF kinases. Consequently, inhibitors designed for cancer can have unpredictable, toxic effects on other organs. Effects on the heart are of particular concern, and there are increasing reports of cancer therapy-related cardiac dysfunction (CTRCD) during treatment and in cancer survivors (e.g. arrhythmias, reduced LVEF or heart failure).^13^ Guidelines focus on monitoring patients at risk of CTRCD,^14^ but the mechanisms underlying CTRCD and how individual therapies affect cardiac function remain largely unknown.

Here, we analysed echocardiograms collected from preclinical mouse studies of cardiac hypertrophy induced by AngII (0.8 mg/kg/d), exploiting the data for greater insight into cardiac function. Instead of M-mode (that predicts values from a single plane across the LV), we used speckle-tracking (strain) to obtain global measures of cardiac dimensions and function. Regional longitudinal strain is informative in patients,^15,16^ so we extended the analysis of mouse echocardiograms for segmental strain, demonstrating variation in endocardial movement around the LV. A significant source of variation between studies was the origin of the mice but, within a specific mouse line, AngII had a greater effect in the basal and mid-regions of the LV. Because ERK1/2 signalling is required for cardiac hypertrophy,^11,12,17^ we applied the methods to understand the effects of dabrafenib and/or trametinib (a MEK inhibitor) on cardiac function. Although these inhibitors reduced AngII-induced cardiac hypertrophy, they did not restore cardiac dysfunction. The data demonstrate the value of using speckle-tracking segmental analysis to understand how different drugs affect cardiac function.

## 2. Methods

### 2.1 Ethics statement

Mice were housed at University of Reading or St. George’s University of London BioResource Unit/Facility. Procedures were performed in accordance with UK regulations and the European Parliament Directive 2010/63/EU for animal experiments. Work was undertaken according to local institutional animal care committee procedures and the U.K. Animals (Scientific Procedures) Act 1986 (Project Licence P8BAB0744).

### 2.2 Source of mice and data

The experimental unit was the individual mouse. We used male mice because young female mice do not respond to AngII with the same robust phenotype.^18^ Archive echocardiograms collected for all relevant studies between November 2018 and February 2021 were used. These included previously reported mice,^19-22^ with additional unreported studies (**Supplementary Table S1**). Male C57Bl/6J mice purchased from Charles River U.K. were allowed to acclimatise for two weeks before experimentation. Wild-type mice were bred at University of Reading from mice with heterozygous global deletion of STRN/STRN3.^22^ These were from C57Bl/6N founders, backcrossed for at least 10 generations onto the C57BL/6J background using mice from Charles River (C57Bl/6(R) mice). Housing conditions were as described.^20,23^ Echocardiograms were collected 7 d after initiation of treatments and, for a subset of mice, after 3 d. Speckle-tracking (strain) analysis of all echocardiograms was performed *de novo* for this study. Mouse body weights are in **Supplementary Table S2**.

### 2.3 *In vivo* mouse studies

Studies were conducted in groups of up to 16 mice to ensure collection of high-quality echocardiograms. Potential confounders were minimised by including vehicle and/or AngII treatments in each cage. Mice were randomly allocated to specific groups independently of the individual leading the experiment. No mice were excluded after randomisation. Individuals conducting the studies were not blinded to experimental conditions for welfare monitoring purposes. Animals were checked daily. Mice undergoing procedures were routinely monitored using a score sheet and culled if they reach a predefined endpoint. No other exclusion criteria were set *a priori* for the experimental process. Following anaesthesia, mice were allowed to recover individually and returned to a clean cage with the original group. At the end of the experiment, mice were culled by cervical dislocation with severance of the femoral artery to ensure cessation of life.

Alzet® osmotic pumps (1004 or 1007D) used for drug delivery were filled according to the manufacturer’s instructions and incubated in sterile PBS (37°C, overnight). For inhibitors (Selleck Chemicals), minipumps contained DMSO/PEG [50% (v/v) dimethyl sulphoxide (DMSO), 20% (v/v) polyethylene glycol 400, 5% (v/v) propylene glycol, 0.5% (v/v) Tween 80], 1 mg/kg/d trametinib (a dose with proven anti-tumour activity in mouse xenografts^24,25^) or 3 mg/kg/d dabrafenib (related to patient dosage of 300 mg/d^26^ https://brimr.org/protein-kinase-inhibitors/) dissolved in DMSO/PEG, or trametinib with dabrafenib. A second minipump was implanted containing acidified PBS or AngII (Merck) in acidified PBS. Buprenorphine was administered subcutaneously for analgesia (Ceva Animal Health Ltd.; 0.05 mg/kg in PBS). Mice were anaesthetised in an induction chamber (vaporised 5% isoflurane with 2 l/min oxygen) and placed on a heated mat in the prone position. Anaesthesia was maintained using a nose cone (2.5% isoflurane in 2 l/min oxygen). The fur covering the mid-scapular region was removed using an electric razor and the area sterilised with HIBISCRUB^®^ (VioVet). A 2 cm incision was made at the mid-scapular region, and blunt dissection generated a pocket towards the lower-left flank of the mouse for minipump insertion. The wound was closed with two simple interrupted sutures using polypropylene 4-0 thread (Prolene, Ethicon).

### 2.4 Mouse echocardiography and analysis of echocardiograms

Echocardiography was performed with a Vevo 2100^TM^ (Fujifilm Visualsonics) using a 38 MHz MS400 transducer. Anaesthesia was induced and maintained using vaporised isoflurane (5% or 1.5%, respectively) in 1 l/min oxygen. Mice were placed on a heated physiological monitoring stage in a supine position; heart rate, respiration rate and body temperature were monitored. Chest fur was removed, pre-warmed ultrasound gel was applied to the chest and the transducer lowered into the gel until a clear image was obtained. Imaging was completed within 10-15 min. Speckle tracking (strain) analysis used B-mode images of long axis views of the heart. VevoStrain software (Fujifilm Visualsonics) was used for analysis. Analysis was performed across two cardiac cycles (heartbeats) starting from the R peak after a breath had been taken (noted by the associated movement of the heart). This provided the most accurate method for tracking endocardial wall movement (a third cycle was often disrupted because of the breathing movement). Global strain measurements and endocardial data for each of 6 segments around the LV were exported. For the latter, measurements (derived from 9 individual points, averaged across the two heartbeats) were collected for radial and longitudinal peak displacement, velocity, strain and strain rate. Images from VevoStrain software were exported as .jpg files and cropped for presentation with Adobe Photoshop CC, maintaining the original relative proportions.

Apart from analysis of individual heartbeats (below), Microsoft Excel and GraphPad Prism 11.0.2 were used for statistical analysis and data presentation. Each echocardiogram was treated independently with reference only to the duration of treatment. For segmental analysis, each segment was treated independently of the others. A Rout outliers test (Q=1%) was used (decided *a priori*) removing a maximum of one outlier for global measurements or two outliers for segmental analysis (<10%). The majority of conditions passed Shapiro-Wilks’ normality tests, so parametric tests were adopted. Two-tailed two-way ANOVA was used with Tukey multiple comparison post-tests to compare C57Bl/6J and C57Bl/6(R) mice and to assess the effects of inhibitors on the cardiac effects of AngII. Specific p values are provided on scatter plots with significance levels (p<0.05) in bold type. For segmental analysis, heatmaps were generated to represent variation in peak values by adjusting all values for each parameter to the means for all segments from vehicle-treated mice. Heatmaps were generated representing the p values from statistical tests.

### 2.5 Heartbeat analysis

For analysis of individual heartbeats, frame-to-frame analysis of radial and longitudinal displacement data was conducted using values for each segment. The analysis preserved the animal as the experimental unit whilst retaining information contained in the full displacement waveform. The data for the two heartbeats were transformed to a shared normalised time axis from 0 to 1 to allow for variation in heart rate and any increase in heart rate resulting from AngII treatment, and to permit comparison of the curves at the same relative heartbeat phase across mice. Scalar mechanical endpoints were calculated for each animal and segment, with trough-to-peak displacement used as the principal measure of dynamic segmental excursion. Heart rate was included as a covariate in all treatment-response models. Treatment effects were assessed relative to the appropriate vehicle control within each cohort; cohort-dependent, time-dependent, and dabrafenib-dependent treatment effects were tested using explicit interaction terms rather than inferred from differences in within-group significance. For each of C57Bl/6J and C57BL/6(R) mice, mice treated with AngII and vehicle were compared; the treatment coefficient is the heart rate-adjusted AngII-vehicle effect. Between these two cohorts, a difference-of-differences model was used: (C57BL/6(R) AngII - C57BL/6(R) Vehicle) - (C57Bl/6J AngII - C57Bl/6J Vehicle). Multiple comparisons were controlled using Benjamini-Hochberg FDR within prespecified families of related tests. Pointwise curve analysis was used as a secondary approach to localise heartbeat phases contributing to significant mechanical differences. Local curve analysis tested the treatment effect at each normalized heartbeat time point after heart rate adjustment. Benjamini-Hochberg FDR correction was applied across the points of each curve. Shaded curve regions in the figures mark FDR-significant heartbeat phases. This hierarchical strategy avoided frame-level pseudo-replication, accounted for heart rate variation and multiple testing, and distinguished global mechanical effects from localised waveform changes, while allowing said changes to serve as important markers for effects of treatment and mouse line.

## 3. Results

### 3.1 Cardiac effects of short-term treatment of C57Bl/6 mice with AngII

Echocardiography was used to determine the acute effects of a slow pressor dose of AngII on mouse hearts, extending the methodology to assess variation of effect around the LV. Because segmental strain analysis is associated with increased variability,^15^ we maximised the power for analysis by using all available, relevant echocardiograms to compare the responses of commercially-available C57Bl/6J mice with C57Bl/6 mice bred in-house (C57Bl/6(R) mice). Mice were treated with 0.8 mg/kg/d AngII or vehicle over 7 d, using osmotic minipumps for drug delivery (**Figure 1A**). Echocardiograms were analysed using speckle-tracking (strain) of B-mode images of the long axis (LAX). Global strain measurements (**Figure 1A-B; Supplementary Table S3**) demonstrated that estimated LV end diastolic mass (LVEDM) was significantly increased by AngII in both C57Bl/6J and C57Bl/6(R) mice, but the time course differed. Thus, the increase in C57Bl/6(R) mice was greater than C57Bl/6J mice at 3 d whereas, by 7 d, the increase in LVEDM was greater in C57Bl/6J mice (**Figure 1C**). End diastolic volume (EDV) and end systolic volume (ESV) were significantly decreased in both mouse lines at 3 d, but the effect was less at 7 d. With respect to function, AngII increased heart rate at 3 d in C57Bl/6(R) but not C57Bl/6J mice, whilst stroke volume was decreased in both lines, and cardiac output declined only in C57Bl/6J mice (**Figure 1D**). AngII did not significantly affect fractional shortening (FS), ejection fraction (EF) or global longitudinal strain (GLS) in either mouse line (**Figure 1E**), consistent with an adaptive hypertrophic response.

**Figure 1.**
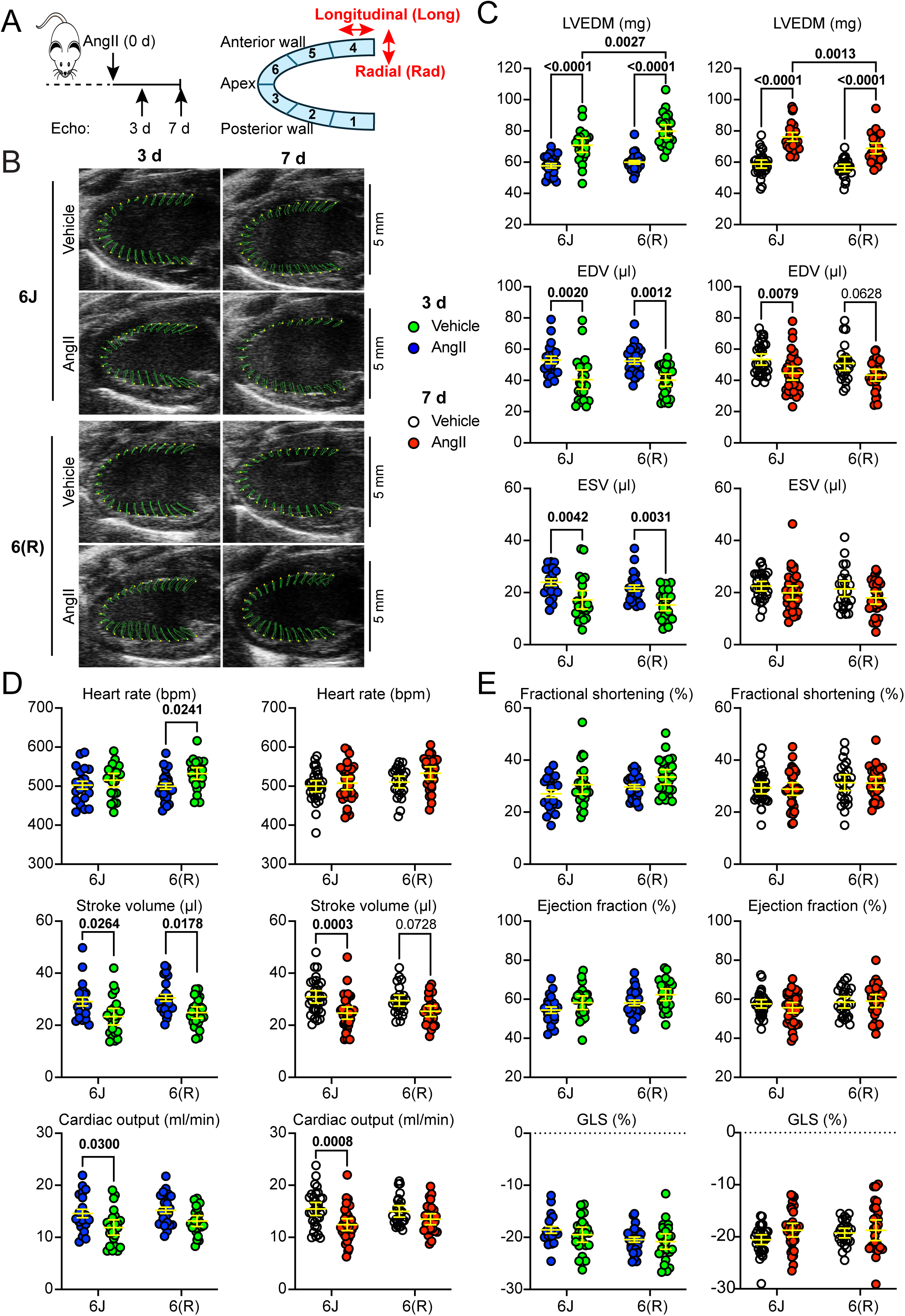
Speckle-tracking (strain) analysis of B-mode images reveals differences in response to AngII in C57Bl/6J compared with C57Bl/6(R) mice. Male mice (10 wks) were treated with 0.8 mg/kg/d AngII or vehicle using osmotic minipumps, and echocardiograms (Echo) collected after 3 or 7 d. **A**, Schematic of experiments (left) and representation of the long axis with segments marked (right). **B**, Representative long axis B-mode images collected at 3 or 7 d from C57Bl/6J (6J) or C57Bl/6(R) (6(R)) mice. **C-E**, Global strain data for left ventricle end diastolic mass (LVEDM), end diastolic volume (EDV) and end systolic volume (ESV) (**C**), heart rate, stroke volume and cardiac output (**D**), and fractional shortening, ejection fraction and global longitudinal strain (GLS) (**E**). Data are from echocardiograms collected at 3 d (left panels; blue/green) and 7 d (right panels; clear/red). Individual points are shown with means ± 95% CI. Statistical analysis used 2-way ANOVA with Tukey’s post-test. Significant differences (p<0.05) are in bold type.

Segmental endocardial speckle-tracking data were generated for echocardiograms from mice treated for 7 d. We assessed peak displacement, velocity, strain and strain rate for longitudinal and radial movement (parallel and perpendicular to the ventricle wall, respectively) of 6 segments around the LV (**Figure 1A**). The effects of AngII in C57Bl/6J mice were most pronounced in the basal region (segments 1 and 4), with a lesser effect in segment 5 (mid-region, anterior wall) (**Figure 2A-C, Supplementary Table S4**). For C57Bl/6(R) mice, no statistically significant differences were detected in peak values in any LV segment in mice treated with AngII *vs* vehicle (**Figure 2A-B and 2D**). For deeper insight, we conducted frame-to-frame analysis of radial and longitudinal displacement, adjusting for heart rate variation. In vehicle-treated C57Bl/6J mouse hearts, LV radial displacement appeared uniform across the two cardiac cycles and, consistent with segmental analysis (**Figure 2**), AngII reduced peak radial displacement only in segment 1 (**Figure 3A**). Longitudinal displacement appeared similar for each cardiac cycle, but whilst the response was uniform in the basal region of the LV, the curves for the mid-regions were flattened, with limited and variable (though consistent between heartbeats) movement at the apex (**Figure 3B**). Overall longitudinal displacement was reduced in segments 1, 2, 4 and 5 in the basal and mid-regions of the LV with AngII treatment (q = 0.012 to <0.001). AngII had a limited effect on trough-to-peak displacement in C57Bl/6(R) mouse hearts, but induced a significant phase shift in radial displacement in the second cardiac cycle despite the heart rate adjustment (**Figure 4A-B**). Assessment of the difference between mouse lines of the differences between vehicle and AngII treatments (difference of differences) showed no significance for radial displacement, but statistical significance was identified in segments 2 and 4 for longitudinal displacement (**Figure 4C-D**).

**Figure 2.**
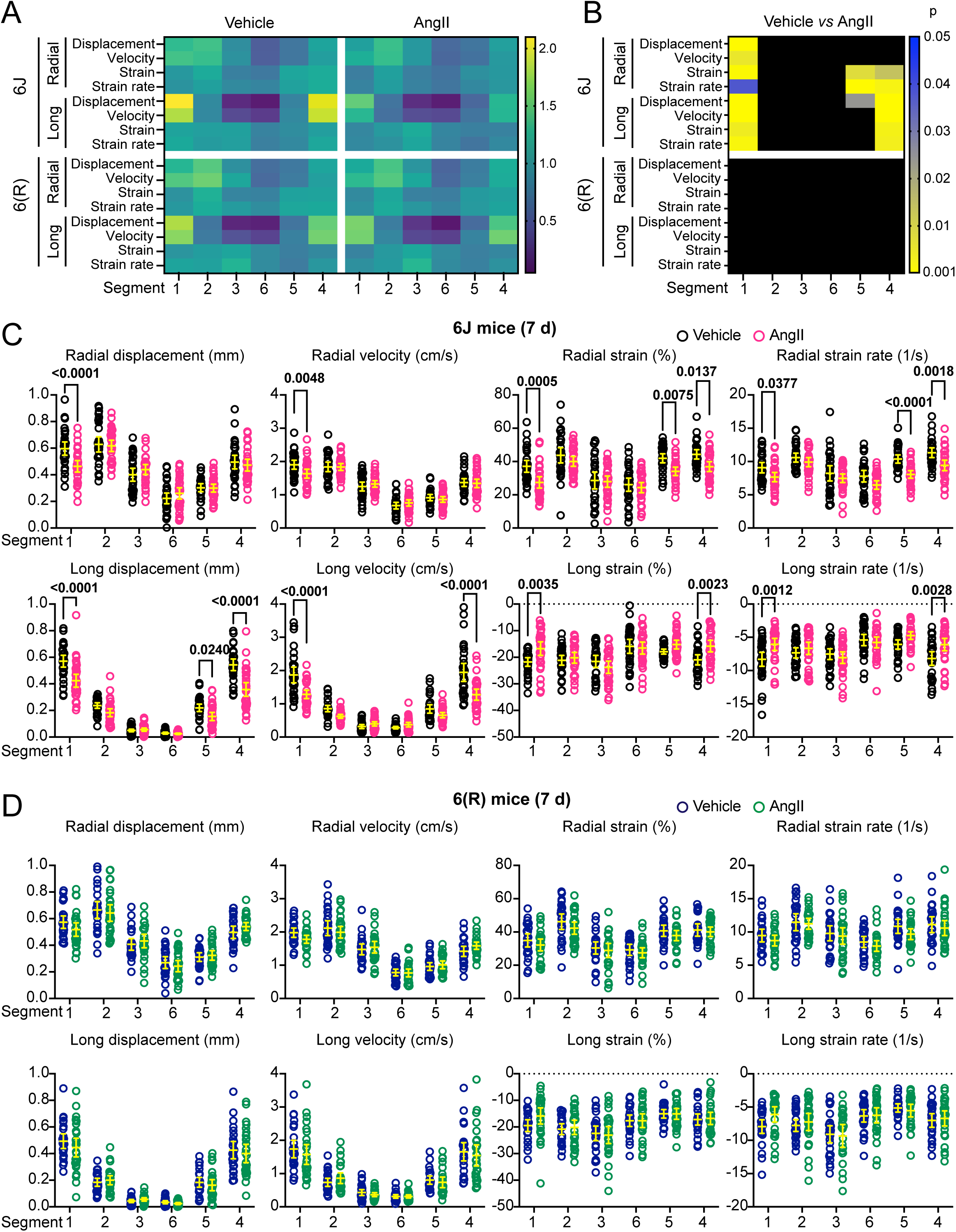
Segmental speckle-tracking demonstrates a predominant effect of AngII on basal and mid sections of the left ventricle in C57Bl/6J mice. Speckle-tracking of long axis B-mode images was performed for 7 d echocardiograms (mice from Figure 1), with export of segmental data. **A**, Heatmaps for each segment around the LV for C57Bl/6J (6J) or C57Bl/6(R) (6(R)) mice showing mean peak values for radial and longitudinal (Long) displacement, velocity, strain and strain rate following treatment with vehicle (left) or AngII (right). Data for each parameter were normalised to the mean value for that parameter in all segments in vehicle-treated C57Bl/6J mice. **B**, Heatmaps for statistical analysis of segmental data showing p values for comparisons between mice treated with vehicle and AngII for 6J and 6(R) mice. **C-D**, Scatter plots showing peak radial (upper graphs) and longitudinal (lower graphs) measurements for each segment for 6J mice treated with vehicle (black) or AngII (pink) (**C**) and 6(R) mice treated with vehicle (blue) or AngII (green) (**D**). Individual values are shown with means ± 95% CI. **B-D**, Analysis used 2-way ANOVA with Tukey’s post-test.

**Figure 3.**
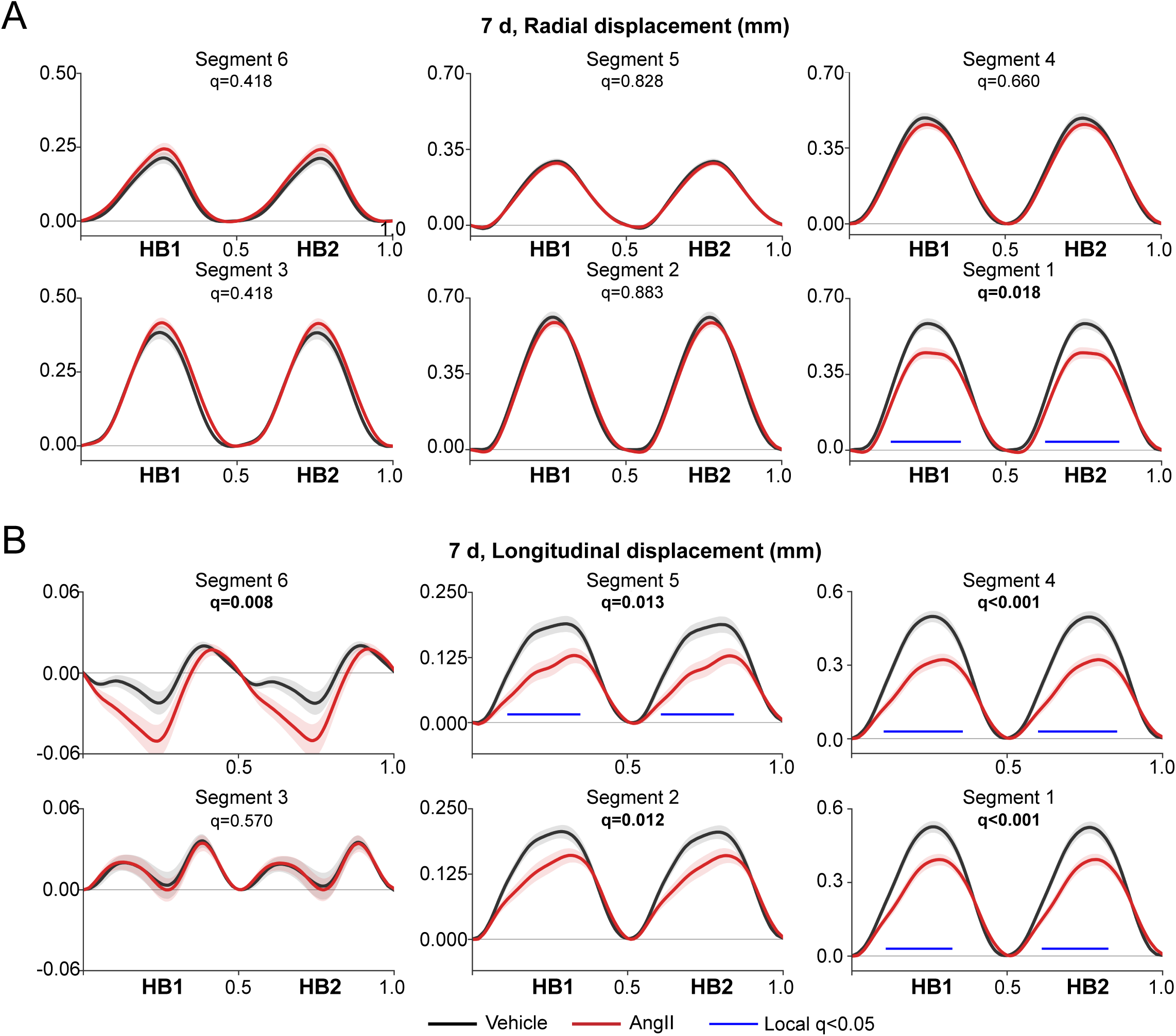
Frame-to-frame analysis of radial and longitudinal displacement identifies effects of AngII in selected segments of the left ventricle in C57Bl/6J mice. Speckle-tracking of long axis B-mode images was performed for 7 d echocardiograms (mice from Figure 1). Radial (**A**) and longitudinal (**B**) displacement data were exported for each frame collected across two heartbeats (HB1, HB2) for each segment around the left ventricle. Data were adjusted for each echocardiogram according to heart rate. Solid lines show mean values, with shaded areas indicating 95% CI. Local statistical significance (q<0.05) is indicated (blue lines) and the overall q value for trough-to-peak values is provided for each segment. Statistically significant values are in bold type.

**Figure 4.**
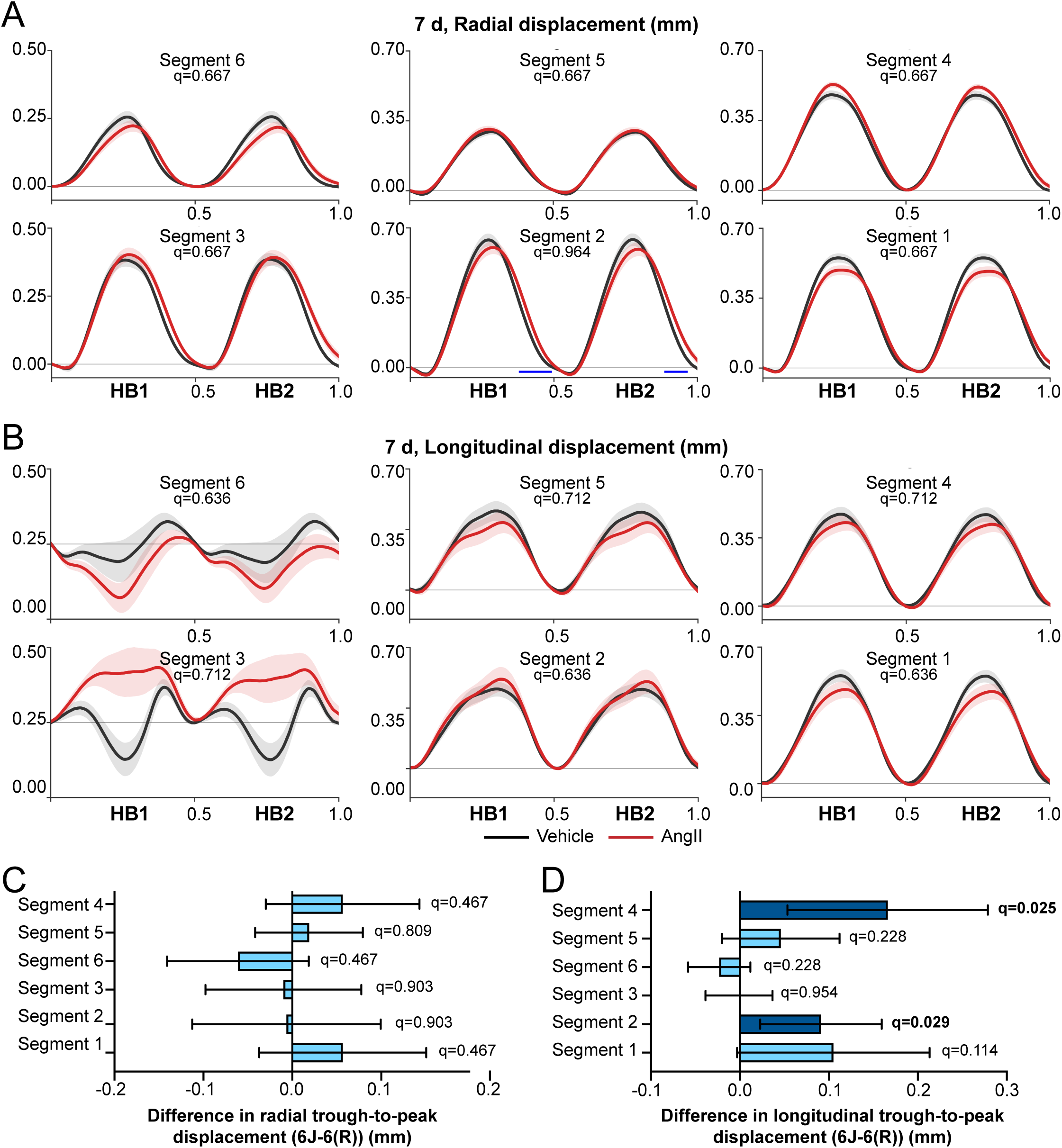
Frame-to-frame analysis of radial and longitudinal displacement identifies limited effects of AngII in selected segments of the left ventricle in C57Bl/6(R) mice. Speckle-tracking of long axis B-mode images was performed for 7 d echocardiograms (mice from Figure 1). Radial (**A**) and longitudinal (**B**) displacement data were exported for each frame collected across two heartbeats (HB1, HB2) for each segment around the left ventricle. Data were adjusted for each echocardiogram according to heart rate. Solid lines show mean values, with shaded areas indicating 95% CI. Local statistical significance (q<0.05) is indicated (blue lines) and the overall q value for trough-to-peak values is provided for each segment. **C-D**, Assessment of the difference of differences between C57Bl/6J and C57Bl/6(R) mice treated with AngII and vehicle for radial (**C**) and longitudinal (**D**) displacement. Statistically significant values are in bold type.

These data demonstrate that 0.8 mg/kg/d AngII promotes early adaptive and rapidly changing hypertrophy with preserved EF in mouse hearts, but the response varies between different mouse lines, a key consideration when comparing different studies. They also illustrate the benefits of using segmental speckle-tracking in preclinical mouse studies. We therefore applied the method to assess the acute effects of dabrafenib and trametinib on cardiac function in C57Bl/6J mice.

### 3.2 Effects of dabrafenib and/or trametinib on cardiac function in C57Bl/6J mice treated without/with AngII

The anti-cancer drugs dabrafenib and trametinib inhibit RAF and MEK kinases, respectively (**Figure 5A**), and may be used individually or together.^12,27^ We previously showed that dabrafenib reduced AngII-induced cardiac hypertrophy over 7 or 28 d, inhibiting cardiac fibrosis, with partial inhibition of cardiomyocyte hypertrophy.^19^ At 28 d, AngII caused cardiac dysfunction with decreased GLS, EF and FS; whilst there was some rescue of GLS and EF by dabrafenib, FS did not recover. Here, we determined if effects of dabrafenib on cardiac function could be detected after just 7 d of treatment using speckle-tracking and segmental analysis of long axis B-mode images. Global strain analysis showed that dabrafenib reduced the AngII-induced increase in LVEDM (**Figure 5B-C, Supplementary Table S5**), consistent with previous M-mode analysis at this time point.^19^ EDV, stroke volume and cardiac output were significantly reduced with AngII treatment (**Figure 5C-D**) (as in the study of all mice, **Figure 1C**). There was no difference in EDV, stroke volume or cardiac output in hearts from mice treated with AngII in the presence of dabrafenib, compared with dabrafenib alone, but this appeared to reflect a reduction with dabrafenib alone rather than, necessarily, normalisation of function (**Figure 5C-D**). None of the treatments significantly affected FS, EF or GLS (**Figure 5E**).

**Figure 5.**
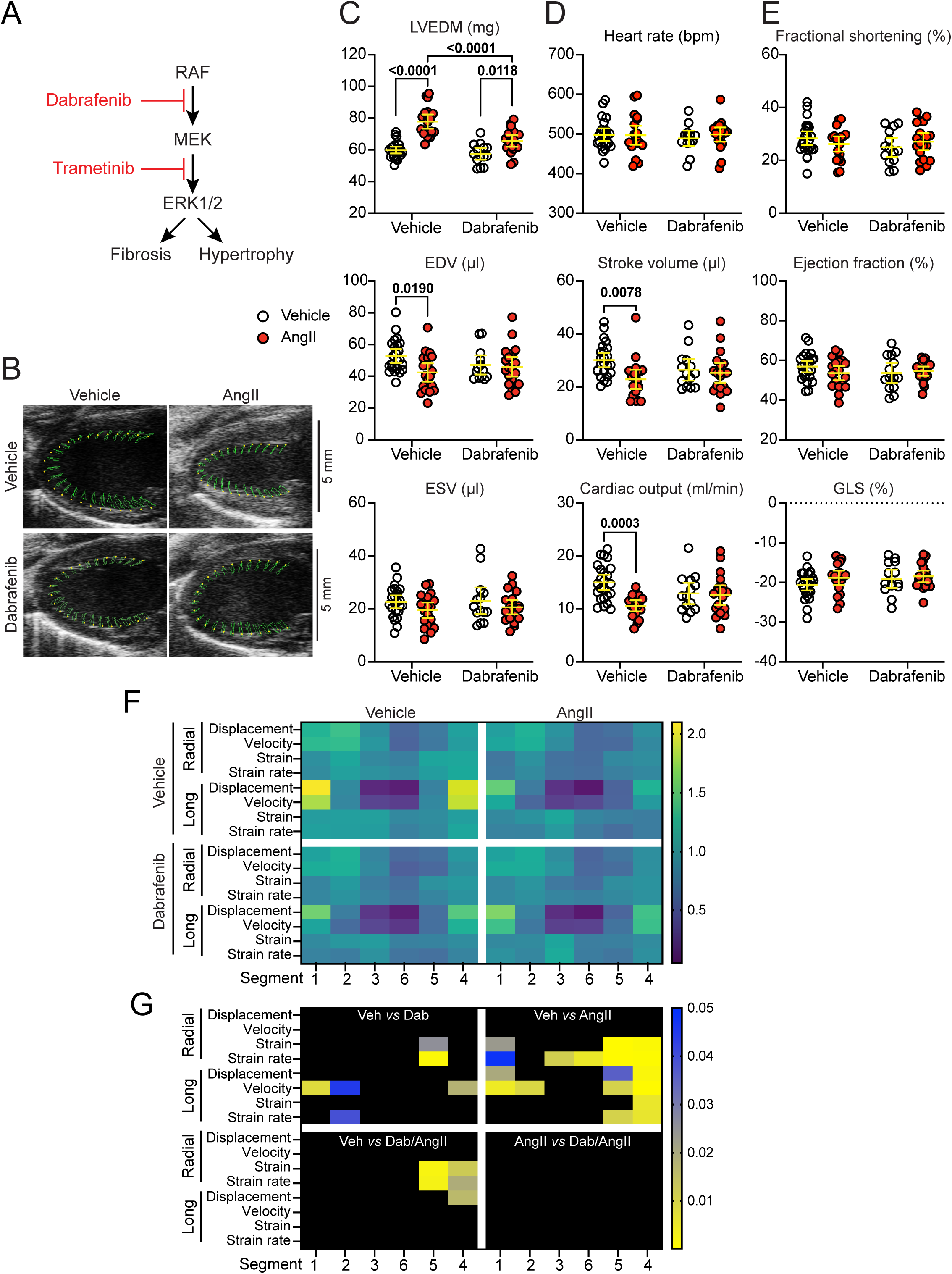
Speckle-tracking of echocardiograms from mice treated with AngII without/with dabrafenib demonstrate that dabrafenib inhibits AngII-induced cardiac hypertrophy but may be detrimental to cardiac function. Male C57Bl/6J mice (10 wks) were treated with vehicle, 3 mg/kg/d dabrafenib (Dab), 0.8 mg/kg/d AngII or AngII with dabrafenib (Dab/AngII) and echocardiograms collected at 7 d. **A**, Diagram of the ERK1/2 cascade and inhibition by dabrafenib and trametinib. **B**, Representative long axis B-mode images showing movement of the endocardial wall across two heartbeats traced in green from the yellow spots. **C-E**, Global strain data for left ventricle end diastolic mass (LVEDM), end diastolic volume (EDV) and end systolic volume (ESV) (**C**), heart rate, stroke volume and cardiac output (**D**), and fractional shortening, ejection fraction and global longitudinal strain (GLS) (**E**). Individual datapoints are shown with means ± 95% CI. **F**, Heatmaps for each segment around the left ventricle showing mean peak values for radial and longitudinal displacement, velocity, strain and strain rate following treatment with vehicle (left) or AngII (right) without (upper) or with (lower) dabrafenib. Data for each parameter were normalised to the mean value for that parameter in all segments in vehicle-treated mice. **G**, Heatmaps for statistical analysis of segmental data showing p values for comparisons between mice treated with the conditions indicated. **C-D** and **G**, Statistical analysis used 2-way ANOVA with Tukey’s post-test.

Segmental strain analysis of LV performance identified similar effects of AngII as in the overall study of C57Bl/6J mice, with the greatest effects in segments 1, 4 and 5, (**Figure 5F-G, Supplementary Figure S1, Supplementary Table S6**). Dabrafenib alone reduced radial strain and strain rate in segment 5, and longitudinal velocity in segments 1 and 4, but it had no significant effects on the changes induced by AngII. Frame-to-frame analysis of longitudinal displacement confirmed that dabrafenib itself reduced trough-to-peak values, and it had no significant effect on the response to AngII (**Figure 6**). Thus, despite apparent normalisation of pathological features of hypertrophy (increase in LVEDM, together with reduced cardiomyocyte hypertrophy and fibrosis^19^), cardiac function was not restored.

**Figure 6.**
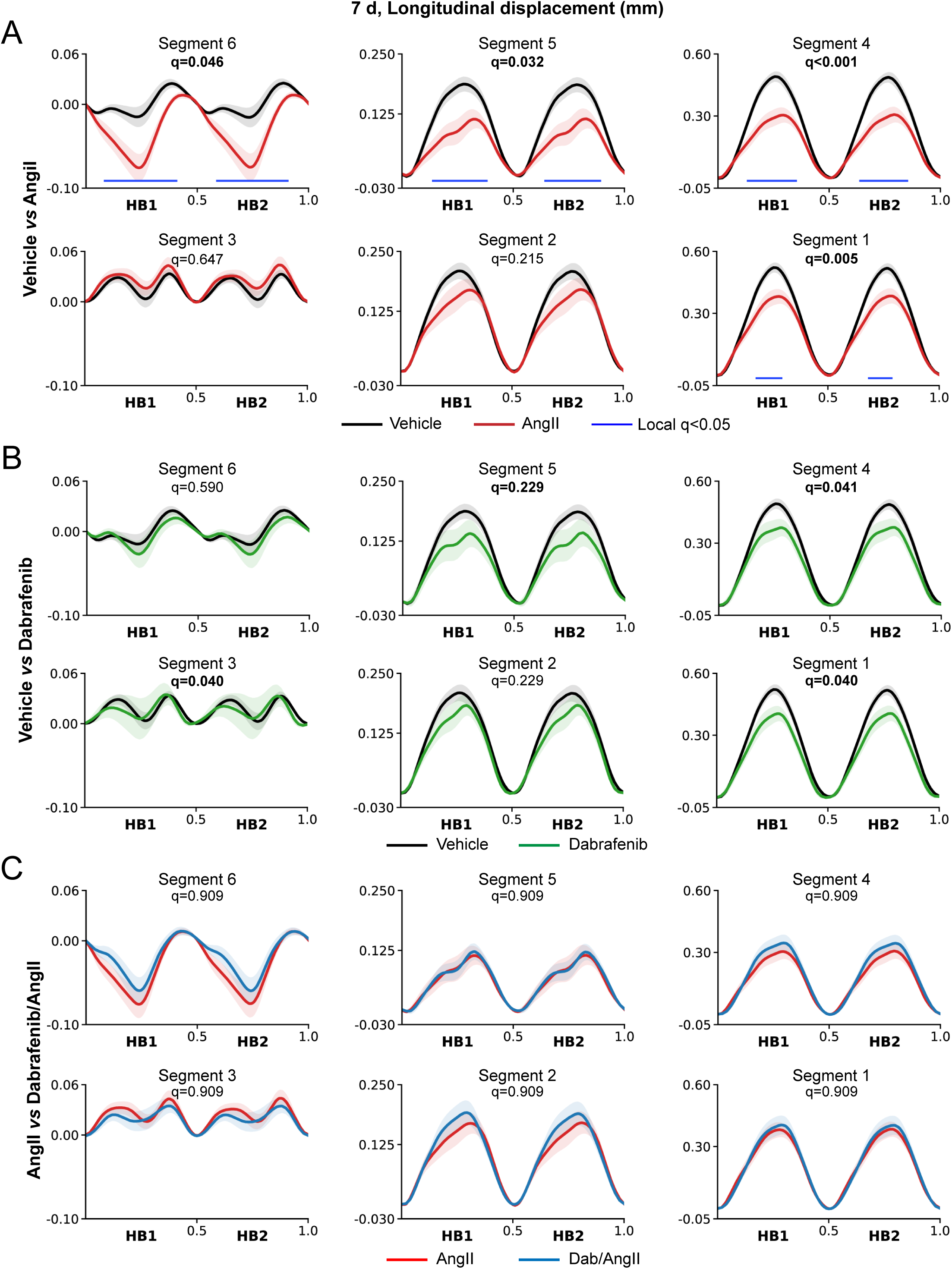
Frame-to-frame analysis of longitudinal displacement for the effects of dabrafenib alone or on the AngII response in segments of the left ventricle in C57Bl/6J mice. Speckle-tracking of long axis B-mode images was performed for 7 d echocardiograms (mice from Figure 5). Longitudinal displacement data were exported for each frame collected across two heartbeats (HB1, HB2) for each segment around the left ventricle. Data were adjusted for each echocardiogram according to heart rate. Solid lines show the mean values, with shaded areas indicating 95% CI. Local statistical significance (q<0.05) is indicated (blue lines) and the overall q value for trough-to-peak measurements is shown for each segment. Statistically significant values are in bold type. Comparisons are shown for vehicle *vs* AngII (**A**), vehicle *vs* dabrafenib (**B**) and AngII *vs* dabrafenib with AngII (**C**).

Trametinib has recognised cardiotoxicity, whether used alone or in combination with dabrafenib.^27^ The effects of trametinib or trametinib/dabrafenib (combination treatment) on AngII-induced hypertrophy over 7 d were assessed in C57Bl/6J mice. The increase in LVEDM induced by AngII was completely inhibited by trametinib or combination treatment (**Figure 7A-B, Supplementary Table S7**). Trametinib or combination treatment alone had no detectable effect on global measures of cardiac function, but caused dysfunction in combination with AngII, with reduced stroke volume and cardiac output, along with EF and GLS (**Figure 7C-D**). Using segmental strain analysis, the effects of trametinib or the combination treatment with AngII compared with vehicle were more pronounced than with AngII alone and confined to the basal and mid-regions of the LV wall (**Figure 7E-F, Supplementary Figure S2, Supplementary Table S8**). Statistically significant differences were not detected between AngII alone and AngII with the inhibitors. The data indicate that trametinib reduced the hypertrophic response to AngII (shown by decreased LVEDM) but increased anomalies in cardiac function resulting from treatment with AngII.

**Figure 7.**
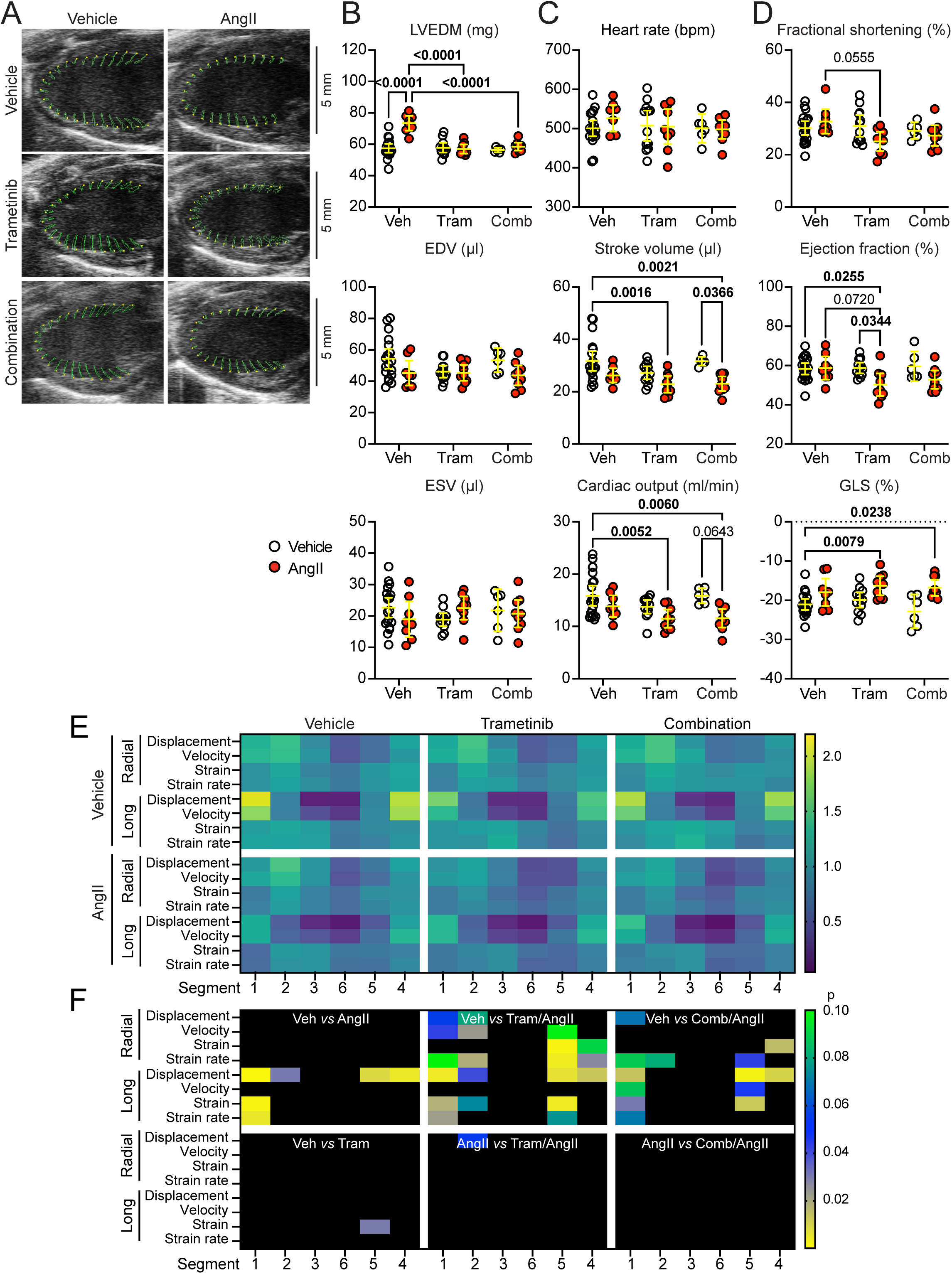
Speckle-tracking of mouse echocardiograms demonstrates that trametinib or the dabrafenib/trametinib combination inhibit AngII-induced cardiac hypertrophy but appear to exacerbate the effect of AngII on endocardial movement. Male C57Bl/6J mice (10 wks) were treated with vehicle, 1 mg /kg/d trametinib (Tram) or trametinib with 3 mg/kg/d dabrafenib (Comb) with/without 0.8 mg/kg/d AngII. Echocardiograms were collected at 7 d. **A**, Representative long axis B-mode images showing the movement of the endocardial wall across two heartbeats traced in green from the yellow spots. **B-D**, Global strain data for left ventricle end diastolic mass (LVEDM), end diastolic volume (EDV) and end systolic volume (ESV) (**B**), heart rate, stroke volume and cardiac output (**C**), and fractional shortening, ejection fraction and global longitudinal strain (GLS) (**D**). Individual datapoints are shown with means ± 95% CI. Statistical analysis used 2-way ANOVA with Tukey’s post-test; significant differences (p<0.05) are in bold type. **E**, Heatmaps for each segment around the left ventricle the showing mean peak values for radial and longitudinal displacement, velocity, strain and strain rate following treatment with vehicle (upper) or AngII (lower) without (left) or with trametinib (centre) or trametinib with dabrafenib (combination; right). Data for each parameter were normalised to the mean value for that parameter in all segments in vehicle-treated mice. **G**, Heatmaps for statistical analysis of segmental data showing p values for comparisons between mice treated with the conditions indicated. Analysis used 2-way ANOVA with Tukey’s post-test.

## 4. Discussion

Much of biomedical sciences research relies on preclinical studies in mice and, for cardiac research, most studies use echocardiography to determine dimensions and assess function. As with patients, mouse echocardiography continues to benefit from improved hardware/software, facilitating collection of more accurate and detailed information. Perhaps disappointingly, reporting usually includes limited data from mouse echocardiograms, often using just M-mode for assessment. For example, standardized echocardiography measures for C57Bl/6N mice reported by the International Mouse Phenotyping Consortium (IMPC) used M-mode imaging of short axis views,^28^ whilst three studies of trametinib in the heart all used M-mode with one reporting only FS and another providing just FS and EF.^29-31^ Here, we demonstrate the strength of speckle-tracking (strain) analysis of mouse 2D echocardiograms which not only provides global measures of dimensions and function but also permits exploration of segmental variation of effects around the LV.

Our studies used C57Bl/6J mice to assess the effects of protein kinase inhibitors (e.g. ^19,20^), but wild-type mice used to assess the roles of STRN/STRN3 in the heart were from C57Bl/6N founders bred in-house against C57Bl/6J mice.^22^ The hypertrophic response induced by AngII over 7 d in wild-type mice from STRN/STRN3 colonies seemed different from commercial C57Bl/6J mice, an observation confirmed here using speckle-tracking and segmental strain (**Figures 1-3**). It is well-established that effects of cardiac interventions in preclinical studies are influenced by the genetic background of mice, and there is variation in response even in sub-strains of the C57Bl/6 line. For example, C57Bl/6N mice respond more severely to transverse aortic constriction than C57Bl/6J mice,^32^ and with additional differences in vascular function (e.g. reduced systolic blood pressure in C57Bl/6N mice) that may affect the cardiac response.^33^ An underlying cause may be reduced mitochondrial-derived oxidative stress in C57Bl/6J mice due to loss of nicotinamide nucleotide transhydrogenase (NNT) from the inner mitochondrial membrane.^34,35^ In our studies, extensive backcrossing with C57Bl/6J mice may still have been insufficient to fully remove the influence of the C57Bl/6N background. Another factor could be breeding and housing conditions in different facilities. Irrespective of the reason, the results add to the body of data that highlight the differences between lines of mice from different facilities.

In all preclinical studies, it is important to maximise the information gained from each experiment. This study demonstrates that greater insight can be gained from extending analysis of mouse echocardiograms beyond M-mode. Clinical studies consider GLS as an earlier marker of cardiac dysfunction than EF, and GLS is used to assess CTRCD in cancer patients.^14^ Segmental/regional variation in longitudinal strain is also important in diseases such as amyloidosis and hypertrophic cardiomyopathy, whilst other cardiac disorders may also exhibit variation in segmental strain.^36-38^ The small size of mouse hearts and high heart rates are challenging but, with high resolution and frame rates, speckle-tracking is possible. One difficulty with segmental analysis is increased variability resulting from fewer datapoints for strain calculation. This was mitigated by maximising group numbers but, although powerful data were obtained with group sizes >20, smaller group sizes still provided useful information. Thus, a significant effect of dabrafenib on endocardial wall movement was detected using segmental strain, with the effect in the basal and mid-regions of the LV (**Figure 5**) and, because the effect of trametinib with AngII was more pronounced than AngII alone, we could detect effects of the inhibitor with AngII not apparent with AngII alone even with smaller groups (**Figure 6**). Frame-to-frame analysis of displacement across two heartbeats allowed more detailed visualisation of endocardial wall movement (**Figures 3-4**). Although variation between heartbeats was largely eliminated by adjusting for heart rate, some displacement of the second heartbeat was detected for radial displacement in segment 2 for C57Bl/6(R) mice. Extending this methodology across more heartbeats would potentially allow detailed assessment of heart rate variability.

Our research focuses on the cardiac effects of protein kinase inhibitors used for cancer, a key pathway being the ERK1/2 cascade.^12^ Of the few published studies of the effects of RAF/MEK inhibitors on mouse hearts, the emphasis has been trametinib for which there are three previously published studies.^29-31^ Beck et al.^29^ identified severe cardiotoxicity with trametinib causing significant reductions in EF and FS, reduced survival rate, increased oxidative stress and marked changes in the transcriptome. Fisher-Wellman et al.^30^ determined that trametinib causes mitochondrial damage even within 3-14 d, with activation of the innate immune response. This could account for the oxidative stress seen in Beck et al.,^29^ and could link to the third study from Fu et al.^31^ who demonstrated an increase in HMGB1, a DAMP (damage-associated molecular pattern) that triggers the immune response.^39^ All studies identified a decline in FS, whilst Fu et al. also reported decreases in EF and cardiac output; none identified any changes in gross morphology with trametinib. We also did not detect any changes in gross morphology in mice treated with trametinib alone (data not shown), but neither did we detect any significant anomalies in mouse hearts treated with trametinib alone using echocardiography (**Figure 7**). The studies have major differences in experimental approach: one used C57Bl/6 mice of both sexes with trametinib delivered in food (1-1.5 mg/kg/d estimated average dose; 120 days)^29^, another treated female FVB mice by oral gavage (3 mg/kg/d; 14 d)^30^, the third treated C57Bl/6J mice (undefined sex) by oral gavage (1.2 mg/kg/d; 7 weeks)^31^, and, in our study, male C57Bl/6J mice received 1 mg/kg/d using osmotic minipumps. FBV/N mice have altered mitochondrial metabolism compared with C57Bl/6J mice^40^ which may be a factor in the mitochondrial involvement in these studies.^30^ However, there is great variation in dose/mode of delivery of AngII, in addition to sex, source, genetic background and origin of the mice, all of which may contribute to the differences in response.

The variation in drug effects between preclinical studies leads to questions about experimental reproducibility and validity of conclusions, if not the entire relevance of conducting such studies using inbred lines. However, inbred lines minimise genetic variation within a study, reducing the numbers of animals required to obtain a meaningful response. Minimally, they provide a starting point to understand potential consequences of drug treatments at least for some patients; at best, they can provide insight into the mechanistic basis of disease. The lack of genetic variation can be mitigated by comparing effects in mice with different background strains as shown with the Mouse Collaborative Cross, a panel of genetically diverse mouse strains used for complex trait analysis.^41^ These mice were used to demonstrate the variation in response to myocardial infarction resulting from genetic background and enabled identification of quantitative trait loci (QTL).^42^ Whilst a full Mouse Collaborative Cross panel would not be possible for all studies, focusing on 2 or 3 lines (rather than just one) may be useful. The challenge is to select appropriate mouse lines for study. For example, an 8-mouse inter-cross study for blood pressure QTL, reported average systolic blood pressures between ∼100 and ∼130 mmHg in different lines^43^ so, for blood pressure studies it may be appropriate to use mouse lines from either end of the spectrum, from the median or with specific a QTL. We suggest that it is not only important to accept the validity of the variation in responses in mice with different genetic backgrounds (as in responses in different patients), but perhaps it is important to capitalise upon it.

Dabrafenib is one of three inhibitors approved for cancers driven by BRAF(V600E) mutations.^44^ Because some inhibitors paradoxically activate ERK1/2 at low concentrations and tumours become resistant to them, they are generally used with a MEK inhibitor (e.g. dabrafenib with trametinib^12^). Some cardiotoxicities were identified with RAF inhibitors, but trametinib (alone or with dabrafenib) has more reported cardiac adverse events including decreased EF, pulmonary embolism (and other thrombotic events), hypertension, atrial fibrillation, cardiac failure, and Qt prolongation.^45^ It is difficult to relate these observations to preclinical studies of trametinib in mice. All the studies used supraphysiological concentrations of trametinib relative to the recommended patient dose (2 mg/d orally for adults >50 kg, maximumally 0.04 mg/kg/d), but within the range used in preclinical studies of cancer (as reported in ^29^). There is also consideration of the variation in response of mice with different genetic backgrounds as discussed above, and, possibly most importantly, none of the studies allow for any systemic effects of cancer. Further refinement in model development and analysis of cardiac function is clearly necessary. Extending the methodology and reporting for segmental strain is one way forward.

In contrast to earlier reports of the effects of trametinib,^29-31^ our studies of dabrafenib (at a patient-relevant dose) and/or trametinib alone showed no significant effect on global measures of cardiac function (^19^; **Figures 5D-E** and 7C-D), although dabrafenib (but not trametinib or the combination) appeared to have some detrimental effects in some LV segments (**Figures 5F-G** and **6**). AngII not only increases blood pressure but also directly affects cardiac cells via the ERK1/2 cascade to promote cardiomyocyte hypertrophy and cardiac fibrosis,^46-48^ so we surmised that RAF/MEK inhibitors would have a more significant effect in model of AngII-induced cardiac hypertrophy. Accordingly, dabrafenib, trametinib or the combination inhibited cardiac hypertrophy induced by AngII (^19^; **Figures 5-7**). However, cardiac function was not restored. There are two main contributing factors. First, the consequence of inhibiting cardiomyocyte hypertrophy without relieving ongoing stress is that the increased workload is not matched by increased cardiomyocyte contractile ability. Secondly, ERK1/2 signalling affects cardiomyocyte contractility directly, for example, by modulating the sodium proton exchanger NHE1,^49^ and inhibiting ERK1/2 is likely to cause dysregulation of ion fluxes between and within cells. It was notable that the effects of AngII and inhibitors were largely on endocardial movement in the basal and mid-regions of the LV (**Figures 5-7**), possibly because of the physical constraints at the apex that limit longitudinal movement. Increasing the number of datapoints per segment would potentially improve imaging of mouse echocardiograms, particularly in this region. With new technological developments, 3D imaging of mouse hearts could offer significantly greater insight into the cardiac effects of cancer drugs with additional assessment of torsion and circumferential strain in different segments.

In conclusion, echocardiography is an important tool in preclinical studies of the heart, but it is rarely used to full potential and reporting of data is often selective. This study demonstrates how speckle-tracking and segmental strain analysis can be used to dissect the effects of different drugs on the LV, extending the methodologies for assessing cardiac function even from archive mouse echocardiograms. Our studies of dabrafenib and trametinib particularly highlight the importance of extending assessment of cardiac function of mouse hearts, not just dimensions. Further advances in hardware/software will continue to increase the accuracy of such measurements and extend the options. From a translational perspective, the data raise the possibility of using speckle-tracking with patient echocardiograms for early identification of CTRCD from specific drugs.

## Abbreviations

AngII: angiotensin II
CTRCD: cancer therapy-related cardiac dysfunction
EDV: end diastolic volume
EF: ejection fraction
ESV: end systolic volume
ERK: extracellular signal-regulated kinase
FS: fractional shortening
GLS: global longitudinal strain
LAX: long axis
LV: left ventricle
LVEDM: LV end diastolic mass
QTL: quantitative trait loci

## Data Availability

All data are available from the authors on request.

