## Supplementary Tables and Figures for "Speckle-tracking echocardiography in preclinical mouse models: Implications for cancer therapy-related cardiac dysfunction from inhibition of the extracellular signal-regulated kinase 1/2 (ERK1/2) cascade"

**Supplementary Table S1. Summary of mouse groups and numbers.**

**Supplementary Table S2. Mouse body weights.**

**Supplementary Table S3. Global strain measures for C57Bl/6J *vs* C57Bl6(R) mice.**

**Supplementary Table S4. Segmental strain measures C57Bl/6J vs C57Bl6(R) mice.**

**Supplementary Table S5. Global strain measures for effects of dabrafenib on C57Bl/6J mouse hearts.**

**Supplementary Table S6. Segmental strain measures for effects of dabrafenib on C57Bl/6J mouse hearts.**

**Supplementary Table S7. Global strain measures for effects of trametinib alone or with dabrafenib (combination) on C57Bl/6J mouse hearts.**

**Supplementary Table S8. Segmental strain measures for effects of trametinib alone or with dabrafenib (combination) on C57Bl/6J mouse hearts.**

**Supplementary Figure S1. Segmental strain data for effects of dabrafenib on AngII-induced hypertrophy: scatter plots for peak values**

**Supplementary Figure S2. Segmental strain data for effects of trametinib on AngII-induced hypertrophy: scatter plots for peak values**

**Supplementary Table S1. Summary of mouse groups and numbers.**

### Experiment conducted in parallel with others reported in [1], but not reported.

§ Experiment conducted de novo for this study; these data have not been reported previously.

#### A. Comparison of C57Bl/6J and C57Bl/6(R) mice.

| | C57Bl/6J | | C57Bl/6(R) | | References |
|---|---|---|---|---|---|
| **Group** | **Vehicle** | **AngII** | **Vehicle** | **AngII** | |
| Previously published | 34 | 22 | 26 | 28 | [1-4] |
| Unpublished | 10 | 12 | ---- | ---- | § |
| **TOTAL** | **44** | **34** | **26** | **28** | |

#### B. Assessment of effects of dabrafenib (Dab).

| **Group** | **Vehicle** | **Dab** | **AngII** | **Dab/AngII** | **References** |
|---|---|---|---|---|---|
| Previously published | 21 | 12 | 15 | 15 | [1] |
| Unpublished | 3 | 0 | 4 | 4 | # |
| **TOTAL** | **24** | **12** | **19** | **19** | |

#### C. Assessment of effects of trametinib (Tram) or trametinib with dabrafenib (combination, Comb).

| **Group** | **Vehicle** | **Tram** | **Comb** | **AngII** | **Tram/ AngII** | **Comb/ AngII** | **References** |
|---|---|---|---|---|---|---|---|
| Previously published | 12 | 12 | 6 | 0 | 0 | 0 | [1,2] |
| Unpublished | 7 | 0 | 0 | 8 | 9 | 9 | § |
| **TOTAL** | **19** | **12** | **6** | **8** | **9** | **9** | |

**Supplementary Table S2. Mouse body weights.** BL, baseline. PS, post minipump surgery.

**A. Comparison of C57Bl/6J and C57Bl/6(R) mice**

| | **C57Bl/6J** | | | | **C57Bl/6(R)** | | | |
|---|---|---|---|---|---|---|---|---|
| | **Vehicle (n=44)** | | **AngII (n=34)** | | **Vehicle (n=30)** | | **AngII (n=28)** | |
| | Mean | SD | Mean | SD | Mean | SD | Mean | SD |
| **BL** | 24.78 | 2.26 | 24.78 | 1.99 | 24.51 | 1.90 | 24.53 | 1.53 |
| **PS** | 26.77 | 2.14 | 26.74 | 1.77 | 26.38 | 1.62 | 26.10 | 1.46 |
| **7 d** | 27.80 | 2.09 | 27.32 | 2.03 | 27.04 | 1.61 | 26.54 | 1.55 |

**B. Study of effects of dabrafenib.** Dab, dabrafenib.

| | **Vehicle (n=25)** | | **Dabrafenib (n=14)** | | **AngII (n=19)** | | **Dab/AngII (n=19)** | |
|---|---|---|---|---|---|---|---|---|
| | **Mean** | **SD** | **Mean** | **SD** | **Mean** | **SD** | **Mean** | **SD** |
| **BL** | 25.08 | 2.19 | 25.02 | 2.26 | 25.13 | 2.10 | 24.33 | 1.74 |
| **PS** | 27.18 | 2.11 | 27.21 | 1.68 | 27.17 | 1.63 | 26.09 | 1.49 |
| **7 d** | 28.23 | 2.09 | 27.64 | 1.50 | 27.80 | 2.09 | 26.96 | 2.43 |

**C. Study of effects of trametinib or trametinib plus dabrafenib (combination).** Tram, trametinib. Comb, combination.

| | **Vehicle (n=19)** | | **Trametinib (n=12)** | | **Combination (n=6)** | | **AngII (n=8)** | | **Tram/AngII (n=9)** | | **Comb/AngII (n=9)** | |
|---|---|---|---|---|---|---|---|---|---|---|---|---|
| | **Mean** | **SD** | **Mean** | **SD** | **Mean** | **SD** | **Mean** | **SD** | **Mean** | **SD** | **Mean** | **SD** |
| **BL** | 25.05 | 2.61 | 24.50 | 2.16 | 27.37 | 2.26 | 24.84 | 2.09 | 23.37 | 2.33 | 23.17 | 1.83 |
| **PS** | 26.89 | 2.26 | 26.13 | 2.44 | 29.17 | 1.72 | 26.13 | 2.22 | 25.44 | 2.50 | 25.11 | 1.78 |
| **7 d** | 27.70 | 1.91 | 26.86 | 1.51 | 28.95 | 1.14 | 26.91 | 2.10 | 25.98 | 2.05 | 25.54 | 1.57 |

**Supplementary Table S3. Global strain measurements for C57Bl/6J *vs* C57Bl6(R) mice.** Analysis used long axis views of the left ventricle (LV) of echocardiograms collected at 3 d (**A**) or 7 d (**B**). EDV, end diastolic volume; ESV, end systolic volume; SV, stroke volume; FS, fractional shortening; CO, cardiac output; GLS, global longitudinal strain; EF, ejection fraction; LVEDM, LV end diastolic mass.

| **A. 3 d** | **C57Bl/6J** | | | | | | **C57Bl/6(R)** | | | | | |
|---|---|---|---|---|---|---|---|---|---|---|---|---|
| | **Vehicle** | | | **AngII** | | | **Vehicle** | | | **AngII** | | |
| | **Mean** | **SD** | **N** | **Mean** | **SD** | **N** | **Mean** | **SD** | **N** | **Mean** | **SD** | **N** |
| **EDV (µl)** | 53.05 | 10.63 | 20 | 40.52 | 14.28 | 23 | 52.29 | 9.14 | 24 | 40.14 | 9.49 | 25 |
| **ESV (µl)** | 23.97 | 5.75 | 20 | 17.22 | 8.15 | 23 | 21.74 | 5.62 | 24 | 15.26 | 5.43 | 25 |
| **SV (µl)** | 29.08 | 7.42 | 20 | 23.30 | 7.31 | 23 | 30.55 | 6.38 | 24 | 24.87 | 5.32 | 25 |
| **FS (%)** | 27.06 | 6.48 | 20 | 30.55 | 8.37 | 23 | 29.68 | 4.64 | 24 | 33.55 | 6.91 | 25 |
| **CO (ml/min)** | 14.58 | 3.61 | 20 | 11.90 | 3.41 | 23 | 15.19 | 2.93 | 24 | 13.14 | 2.52 | 25 |
| **GLS (%)** | -18.56 | 2.93 | 20 | -19.49 | 3.37 | 23 | -20.44 | 2.49 | 24 | -20.82 | 3.56 | 25 |
| **EF (%)** | 54.42 | 7.32 | 20 | 58.22 | 8.11 | 23 | 58.10 | 6.87 | 24 | 62.33 | 7.32 | 25 |
| **LVEDM (mg)** | 57.66 | 6.62 | 20 | 70.82 | 10.44 | 23 | 59.94 | 5.57 | 24 | 79.76 | 10.14 | 25 |
| **Heart rate (bpm)** | 502 | 45 | 20 | 515 | 40 | 23 | 500 | 36 | 24 | 532 | 37 | 25 |

| **B. 7 d** | **C57Bl/6J** | | | | | | **C57Bl/6(R)** | | | | | |
|---|---|---|---|---|---|---|---|---|---|---|---|---|
| | **Vehicle** | | | **AngII** | | | **Vehicle** | | | **AngII** | | |
| | **Mean** | **SD** | **N** | **Mean** | **SD** | **N** | **Mean** | **SD** | **N** | **Mean** | **SD** | **N** |
| **EDV (µl)** | 53.38 | 10.00 | 32 | 44.59 | 12.60 | 34 | 50.86 | 11.04 | 26 | 43.35 | 9.69 | 28 |
| **ESV (µl)** | 22.47 | 5.11 | 32 | 19.91 | 7.70 | 34 | 21.44 | 7.47 | 26 | 17.97 | 6.37 | 28 |
| **SV (µl)** | 30.91 | 6.56 | 32 | 24.68 | 6.73 | 34 | 29.42 | 5.19 | 26 | 25.37 | 5.25 | 28 |
| **FS (%)** | 29.30 | 6.16 | 32 | 28.90 | 6.94 | 34 | 31.30 | 7.56 | 26 | 31.19 | 6.21 | 28 |
| **CO (ml/min)** | 15.46 | 3.48 | 32 | 12.44 | 3.25 | 34 | 14.99 | 2.75 | 26 | 13.48 | 2.75 | 28 |
| **GLS (%)** | -20.56 | 2.60 | 32 | -18.73 | 3.75 | 34 | -19.30 | 2.29 | 26 | -18.74 | 5.13 | 28 |
| **EF (%)** | 57.64 | 5.97 | 32 | 55.63 | 7.71 | 34 | 58.40 | 7.17 | 26 | 59.06 | 8.54 | 28 |
| **LVEDM (mg)** | 58.78 | 7.18 | 32 | 76.19 | 8.15 | 34 | 56.30 | 5.81 | 26 | 68.76 | 8.93 | 28 |
| **Heart rate (bpm)** | 500 | 40 | 32 | 507 | 47 | 34 | 511 | 37 | 26 | 533 | 42 | 28 |

**Supplementary Table S4. Segmental strain measurements C57Bl/6J *vs* C57Bl6(R) mice.** Analysis used long axis views of the left ventricle (LV). **A**, C57Bl/6J mice. **B**, C57Bl/6(R) mice. Long, longitudinal.

| A. C57Bl/6J mice | | Vehicle | | | AngII | | |
|---|---|---|---|---|---|---|---|
| | Segment | Mean | SD | Mean | SD | Mean | SD |
| Radial displacement (mm) | 1 | 0.597 | 0.139 | 32 | 0.464 | 0.122 | 33 |
| Radial displacement (mm) | 2 | 0.629 | 0.158 | 32 | 0.618 | 0.121 | 34 |
| Radial displacement (mm) | 3 | 0.400 | 0.135 | 32 | 0.440 | 0.106 | 34 |
| Radial displacement (mm) | 6 | 0.223 | 0.111 | 32 | 0.261 | 0.121 | 34 |
| Radial displacement (mm) | 5 | 0.303 | 0.084 | 32 | 0.300 | 0.085 | 34 |
| Radial displacement (mm) | 4 | 0.496 | 0.141 | 32 | 0.473 | 0.128 | 34 |
| Radial velocity (cm/s) | 1 | 1.903 | 0.373 | 32 | 1.636 | 0.382 | 34 |
| Radial velocity (cm/s) | 2 | 1.839 | 0.396 | 32 | 1.836 | 0.296 | 34 |
| Radial velocity (cm/s) | 3 | 1.249 | 0.366 | 32 | 1.326 | 0.279 | 34 |
| Radial velocity (cm/s) | 6 | 0.674 | 0.273 | 32 | 0.755 | 0.266 | 34 |
| Radial velocity (cm/s) | 5 | 0.915 | 0.258 | 32 | 0.859 | 0.242 | 34 |
| Radial velocity (cm/s) | 4 | 1.368 | 0.314 | 31 | 1.341 | 0.360 | 34 |
| Radial strain (%) | 1 | 37.10 | 9.85 | 32 | 27.46 | 9.90 | 34 |
| Radial strain (%) | 2 | 43.73 | 12.72 | 32 | 40.07 | 8.51 | 34 |
| Radial strain (%) | 3 | 28.57 | 14.00 | 32 | 27.88 | 9.70 | 34 |
| Radial strain (%) | 6 | 26.07 | 10.95 | 32 | 24.29 | 8.72 | 34 |
| Radial strain (%) | 5 | 41.94 | 7.47 | 32 | 34.10 | 7.30 | 34 |
| Radial strain (%) | 4 | 44.31 | 7.77 | 32 | 36.91 | 8.48 | 34 |
| Radial strain rate (1/s) | 1 | 9.08 | 2.18 | 32 | 7.65 | 2.01 | 34 |
| Radial strain rate (1/s) | 2 | 10.58 | 1.86 | 31 | 9.91 | 1.94 | 34 |
| Radial strain rate (1/s) | 3 | 8.23 | 3.20 | 32 | 7.48 | 1.83 | 34 |
| Radial strain rate (1/s) | 6 | 7.79 | 2.41 | 32 | 6.47 | 1.81 | 34 |
| Radial strain rate (1/s) | 5 | 10.37 | 1.72 | 32 | 8.06 | 1.62 | 34 |
| Radial strain rate (1/s) | 4 | 11.33 | 2.01 | 31 | 9.42 | 2.27 | 34 |
| Long displacement (mm) | 1 | 0.571 | 0.123 | 32 | 0.424 | 0.139 | 34 |
| Long displacement (mm) | 2 | 0.237 | 0.057 | 32 | 0.183 | 0.085 | 34 |
| Long displacement (mm) | 3 | 0.050 | 0.027 | 32 | 0.056 | 0.034 | 34 |
| Long displacement (mm) | 6 | 0.029 | 0.021 | 32 | 0.025 | 0.016 | 31 |
| Long displacement (mm) | 5 | 0.220 | 0.077 | 32 | 0.155 | 0.086 | 34 |
| Long displacement (mm) | 4 | 0.543 | 0.121 | 32 | 0.358 | 0.149 | 34 |
| Long velocity (cm/s) | 1 | 1.868 | 0.652 | 32 | 1.284 | 0.360 | 33 |
| Long velocity (cm/s) | 2 | 0.855 | 0.258 | 31 | 0.625 | 0.158 | 33 |
| Long velocity (cm/s) | 3 | 0.317 | 0.161 | 29 | 0.398 | 0.183 | 34 |
| Long velocity (cm/s) | 6 | 0.286 | 0.103 | 32 | 0.374 | 0.193 | 34 |
| Long velocity (cm/s) | 5 | 0.844 | 0.347 | 31 | 0.660 | 0.221 | 33 |
| Long velocity (cm/s) | 4 | 1.945 | 0.749 | 32 | 1.275 | 0.470 | 33 |
| Long strain (%) | 1 | -21.78 | 4.40 | 31 | -16.76 | 7.34 | 34 |
| Long strain (%) | 2 | -21.15 | 5.14 | 32 | -20.20 | 5.81 | 34 |
| Long strain (%) | 3 | -21.56 | 6.22 | 32 | -23.77 | 6.61 | 34 |
| Long strain (%) | 6 | -15.89 | 7.28 | 32 | -16.53 | 6.51 | 34 |
| Long strain (%) | 5 | -17.88 | 2.59 | 32 | -15.22 | 4.70 | 34 |
| Long strain (%) | 4 | -20.97 | 5.31 | 32 | -15.82 | 6.19 | 34 |
| Long strain rate (1/s) | 1 | -8.30 | 2.92 | 32 | -6.06 | 2.61 | 34 |
| Long strain rate (1/s) | 2 | -7.34 | 2.13 | 32 | -6.66 | 2.62 | 34 |
| Long strain rate (1/s) | 3 | -7.53 | 2.24 | 32 | -8.06 | 2.49 | 34 |
| Long strain rate (1/s) | 6 | -5.41 | 2.34 | 32 | -5.77 | 2.32 | 34 |
| Long strain rate (1/s) | 5 | -6.08 | 1.91 | 32 | -4.75 | 1.63 | 34 |
| Long strain rate (1/s) | 4 | -8.19 | 2.86 | 32 | -6.08 | 2.68 | 34 |

| B. C57Bl/6(R) mice | | Vehicle | | | AngII | | |
|---|---|---|---|---|---|---|---|
| | Segment | Mean | SD | Mean | SD | Mean | SD |
| **Radial displacement (mm)** | 1 | 0.574 | 0.116 | 26 | 0.518 | 0.130 | 28 |
| **Radial displacement (mm)** | 2 | 0.665 | 0.159 | 26 | 0.643 | 0.160 | 28 |
| **Radial displacement (mm)** | 3 | 0.407 | 0.108 | 26 | 0.432 | 0.131 | 28 |
| **Radial displacement (mm)** | 6 | 0.270 | 0.110 | 26 | 0.243 | 0.108 | 28 |
| **Radial displacement (mm)** | 5 | 0.311 | 0.081 | 26 | 0.324 | 0.092 | 28 |
| **Radial displacement (mm)** | 4 | 0.497 | 0.115 | 26 | 0.541 | 0.077 | 28 |
| **Radial velocity (cm/s)** | 1 | 1.980 | 0.347 | 26 | 1.780 | 0.306 | 28 |
| **Radial velocity (cm/s)** | 2 | 2.115 | 0.551 | 26 | 1.992 | 0.445 | 28 |
| **Radial velocity (cm/s)** | 3 | 1.477 | 0.392 | 25 | 1.540 | 0.446 | 28 |
| **Radial velocity (cm/s)** | 6 | 0.783 | 0.246 | 26 | 0.769 | 0.302 | 28 |
| **Radial velocity (cm/s)** | 5 | 0.961 | 0.290 | 26 | 1.011 | 0.299 | 28 |
| **Radial velocity (cm/s)** | 4 | 1.417 | 0.377 | 26 | 1.585 | 0.280 | 28 |
| **Radial strain (%)** | 1 | 34.91 | 9.83 | 26 | 32.17 | 9.19 | 28 |
| **Radial strain (%)** | 2 | 46.19 | 11.54 | 26 | 42.28 | 9.44 | 28 |
| **Radial strain (%)** | 3 | 30.38 | 9.07 | 26 | 28.90 | 10.80 | 28 |
| **Radial strain (%)** | 6 | 29.25 | 8.26 | 26 | 27.50 | 7.80 | 28 |
| **Radial strain (%)** | 5 | 40.72 | 8.36 | 26 | 37.85 | 7.93 | 28 |
| **Radial strain (%)** | 4 | 41.19 | 9.18 | 26 | 40.07 | 7.40 | 28 |
| **Radial strain rate (1/s)** | 1 | 9.47 | 2.38 | 26 | 8.71 | 2.06 | 28 |
| **Radial strain rate (1/s)** | 2 | 11.47 | 3.16 | 26 | 11.24 | 2.11 | 28 |
| **Radial strain rate (1/s)** | 3 | 9.80 | 2.80 | 26 | 9.40 | 3.20 | 28 |
| **Radial strain rate (1/s)** | 6 | 8.50 | 2.05 | 26 | 7.89 | 2.08 | 28 |
| **Radial strain rate (1/s)** | 5 | 10.85 | 2.61 | 26 | 9.67 | 2.10 | 28 |
| **Radial strain rate (1/s)** | 4 | 11.10 | 2.95 | 26 | 10.59 | 2.91 | 28 |
| **Long displacement (mm)** | 1 | 0.494 | 0.137 | 26 | 0.446 | 0.180 | 28 |
| **Long displacement (mm)** | 2 | 0.185 | 0.074 | 26 | 0.200 | 0.087 | 28 |
| **Long displacement (mm)** | 3 | 0.044 | 0.029 | 26 | 0.056 | 0.032 | 26 |
| **Long displacement (mm)** | 6 | 0.035 | 0.029 | 25 | 0.024 | 0.019 | 28 |
| **Long displacement (mm)** | 5 | 0.184 | 0.092 | 26 | 0.163 | 0.108 | 28 |
| **Long displacement (mm)** | 4 | 0.426 | 0.161 | 26 | 0.401 | 0.178 | 28 |
| **Long velocity (cm/s)** | 1 | 1.737 | 0.654 | 26 | 1.574 | 0.751 | 27 |
| **Long velocity (cm/s)** | 2 | 0.737 | 0.299 | 26 | 0.861 | 0.409 | 28 |
| **Long velocity (cm/s)** | 3 | 0.431 | 0.219 | 26 | 0.367 | 0.153 | 28 |
| **Long velocity (cm/s)** | 6 | 0.313 | 0.134 | 25 | 0.312 | 0.125 | 28 |
| **Long velocity (cm/s)** | 5 | 0.803 | 0.311 | 26 | 0.737 | 0.394 | 26 |
| **Long velocity (cm/s)** | 4 | 1.656 | 0.683 | 26 | 1.546 | 0.800 | 27 |
| **Long strain (%)** | 1 | -19.48 | 5.46 | 26 | -15.85 | 7.85 | 28 |
| **Long strain (%)** | 2 | -20.81 | 5.03 | 26 | -20.13 | 6.93 | 28 |
| **Long strain (%)** | 3 | -22.40 | 6.78 | 26 | -23.04 | 8.26 | 28 |
| **Long strain (%)** | 6 | -17.65 | 5.36 | 26 | -17.63 | 6.63 | 28 |
| **Long strain (%)** | 5 | -15.03 | 3.95 | 26 | -15.08 | 5.17 | 28 |
| **Long strain (%)** | 4 | -17.20 | 5.18 | 26 | -16.78 | 5.91 | 28 |
| **Long strain rate (1/s)** | 1 | -7.94 | 2.55 | 26 | -6.11 | 2.90 | 27 |
| **Long strain rate (1/s)** | 2 | -7.77 | 2.19 | 26 | -7.24 | 3.30 | 28 |
| **Long strain rate (1/s)** | 3 | -9.10 | 3.18 | 26 | -9.34 | 4.43 | 28 |
| **Long strain rate (1/s)** | 6 | -6.32 | 2.31 | 26 | -6.24 | 2.93 | 28 |
| **Long strain rate (1/s)** | 5 | -5.14 | 1.54 | 26 | -5.54 | 2.46 | 28 |
| **Long strain rate (1/s)** | 4 | -7.01 | 2.54 | 26 | -6.71 | 2.93 | 28 |

**Supplementary Table S5. Global strain measurements for effects of dabrafenib on C57Bl/6J mouse hearts.** Analysis used long axis views of the left ventricle (LV) of echocardiograms collected at 7 d. EDV, end diastolic volume; ESV, end systolic volume; SV, stroke volume; FS, fractional shortening; CO, cardiac output; GLS, global longitudinal strain; EF, ejection fraction; LVEDM, LV end diastolic mass.

| | **Vehicle** | | | **AngII** | | |
|---|---|---|---|---|---|---|
| | **Mean** | **SD** | **N** | **Mean** | **SD** | **N** |
| **EDV (µl)** | 52.79 | 10.21 | 24 | 42.38 | 11.95 | 19 |
| **ESV (µl)** | 22.69 | 5.81 | 24 | 19.55 | 6.11 | 19 |
| **SV (µl)** | 30.10 | 6.40 | 24 | 22.82 | 7.55 | 19 |
| **FS (%)** | 28.35 | 6.27 | 24 | 26.19 | 5.99 | 19 |
| **CO (ml/min)** | 15.02 | 3.42 | 24 | 10.59 | 2.11 | 18 |
| **GLS (%)** | -20.56 | 3.45 | 24 | -18.81 | 3.80 | 19 |
| **EF (%)** | 56.89 | 6.98 | 24 | 53.57 | 7.41 | 19 |
| **LVEDM (mg)** | 59.85 | 5.02 | 23 | 77.85 | 9.27 | 19 |
| **Heart rate (bpm)** | 498 | 37 | 24 | 497 | 50 | 19 |
| | **Dabrafenib** | | | **Dabrafenib/AngII** | | |
| | **Mean** | **SD** | **N** | **Mean** | **SD** | **N** |
| **EDV (µl)** | 47.15 | 10.03 | 13 | 45.99 | 12.67 | 19 |
| **ESV (µl)** | 23.04 | 8.70 | 14 | 20.65 | 5.58 | 19 |
| **SV (µl)** | 26.46 | 7.10 | 14 | 25.34 | 7.73 | 19 |
| **FS (%)** | 24.99 | 6.24 | 14 | 27.14 | 6.65 | 19 |
| **CO (ml/min)** | 12.93 | 3.52 | 14 | 12.61 | 3.83 | 19 |
| **GLS (%)** | -19.06 | 3.99 | 14 | -18.46 | 3.07 | 19 |
| **EF (%)** | 53.72 | 8.65 | 14 | 54.59 | 4.86 | 19 |
| **LVEDM (mg)** | 57.55 | 6.54 | 14 | 65.57 | 7.49 | 19 |
| **Heart rate (bpm)** | 489 | 34 | 14 | 499 | 37 | 19 |

**Supplementary Table S6. Segmental strain measurements for effects of dabrafenib on C57Bl/6J mouse hearts.** Analysis used long-axis views of the left ventricle (LV). **A**, Radial measurements. **B**, Longitudinal (Long) measurements.

| A. Radial measurements | | Vehicle | | | AngII | | |
|---|---|---|---|---|---|---|---|
| | Segment | Mean | SD | Mean | SD | Mean | SD |
| Radial displacement (mm) | 1 | 0.573 | 0.148 | 24 | 0.475 | 0.185 | 19 |
| Radial displacement (mm) | 2 | 0.632 | 0.177 | 24 | 0.569 | 0.110 | 19 |
| Radial displacement (mm) | 3 | 0.408 | 0.122 | 24 | 0.431 | 0.111 | 19 |
| Radial displacement (mm) | 6 | 0.254 | 0.126 | 24 | 0.262 | 0.114 | 19 |
| Radial displacement (mm) | 5 | 0.324 | 0.087 | 24 | 0.285 | 0.072 | 19 |
| Radial displacement (mm) | 4 | 0.492 | 0.140 | 24 | 0.412 | 0.108 | 19 |
| Radial velocity (cm/s) | 1 | 1.878 | 0.436 | 24 | 1.591 | 0.435 | 19 |
| Radial velocity (cm/s) | 2 | 1.908 | 0.516 | 24 | 1.729 | 0.275 | 19 |
| Radial velocity (cm/s) | 3 | 1.383 | 0.451 | 24 | 1.247 | 0.237 | 19 |
| Radial velocity (cm/s) | 6 | 0.762 | 0.300 | 24 | 0.811 | 0.225 | 19 |
| Radial velocity (cm/s) | 5 | 0.978 | 0.299 | 24 | 0.826 | 0.205 | 19 |
| Radial velocity (cm/s) | 4 | 1.447 | 0.428 | 24 | 1.198 | 0.342 | 19 |
| Radial strain (%) | 1 | 34.82 | 9.29 | 24 | 25.98 | 10.34 | 19 |
| Radial strain (%) | 2 | 43.42 | 13.45 | 24 | 36.11 | 6.03 | 19 |
| Radial strain (%) | 3 | 32.71 | 14.12 | 24 | 24.70 | 9.72 | 19 |
| Radial strain (%) | 6 | 29.58 | 13.15 | 24 | 22.86 | 8.12 | 19 |
| Radial strain (%) | 5 | 42.81 | 6.56 | 24 | 32.44 | 6.57 | 19 |
| Radial strain (%) | 4 | 43.84 | 7.72 | 24 | 34.36 | 7.03 | 19 |
| Radial strain rate (1/s) | 1 | 9.00 | 1.96 | 24 | 7.17 | 2.05 | 19 |
| Radial strain rate (1/s) | 2 | 10.80 | 3.18 | 24 | 8.87 | 1.60 | 19 |
| Radial strain rate (1/s) | 3 | 9.60 | 3.87 | 24 | 6.71 | 1.83 | 19 |
| Radial strain rate (1/s) | 6 | 8.68 | 2.38 | 24 | 6.32 | 1.58 | 19 |
| Radial strain rate (1/s) | 5 | 10.60 | 1.80 | 24 | 7.73 | 1.47 | 19 |
| Radial strain rate (1/s) | 4 | 11.81 | 2.83 | 24 | 8.58 | 2.04 | 19 |
| | | Dabrafenib | | | Dabrafenib/AngII | | |
| | Segment | Mean | SD | Mean | SD | Mean | SD |
| Radial displacement (mm) | 1 | 0.495 | 0.165 | 13 | 0.496 | 0.126 | 19 |
| Radial displacement (mm) | 2 | 0.560 | 0.170 | 13 | 0.541 | 0.144 | 19 |
| Radial displacement (mm) | 3 | 0.371 | 0.128 | 13 | 0.401 | 0.130 | 19 |
| Radial displacement (mm) | 6 | 0.265 | 0.077 | 13 | 0.246 | 0.106 | 19 |
| Radial displacement (mm) | 5 | 0.288 | 0.055 | 13 | 0.303 | 0.068 | 18 |
| Radial displacement (mm) | 4 | 0.447 | 0.137 | 13 | 0.408 | 0.101 | 18 |
| Radial velocity (cm/s) | 1 | 1.698 | 0.500 | 13 | 1.704 | 0.395 | 19 |
| Radial velocity (cm/s) | 2 | 1.733 | 0.479 | 13 | 1.660 | 0.386 | 19 |
| Radial velocity (cm/s) | 3 | 1.218 | 0.324 | 13 | 1.364 | 0.321 | 19 |
| Radial velocity (cm/s) | 6 | 0.716 | 0.166 | 13 | 0.807 | 0.352 | 19 |
| Radial velocity (cm/s) | 5 | 0.805 | 0.174 | 13 | 0.845 | 0.294 | 19 |
| Radial velocity (cm/s) | 4 | 1.275 | 0.373 | 13 | 1.280 | 0.365 | 19 |
| Radial strain (%) | 1 | 31.31 | 11.36 | 13 | 31.73 | 8.80 | 19 |
| Radial strain (%) | 2 | 38.81 | 11.96 | 13 | 36.93 | 10.48 | 19 |
| Radial strain (%) | 3 | 27.95 | 14.27 | 13 | 26.69 | 10.47 | 19 |
| Radial strain (%) | 6 | 24.07 | 10.90 | 13 | 23.93 | 10.55 | 19 |
| Radial strain (%) | 5 | 35.52 | 8.03 | 13 | 34.23 | 8.35 | 19 |
| Radial strain (%) | 4 | 38.33 | 7.21 | 13 | 36.53 | 7.92 | 19 |
| Radial strain rate (1/s) | 1 | 8.46 | 3.14 | 13 | 8.63 | 2.09 | 19 |
| Radial strain rate (1/s) | 2 | 9.70 | 2.90 | 13 | 9.34 | 2.18 | 19 |
| Radial strain rate (1/s) | 3 | 8.12 | 2.75 | 13 | 8.36 | 2.54 | 18 |
| Radial strain rate (1/s) | 6 | 6.78 | 2.50 | 13 | 7.85 | 2.31 | 19 |
| Radial strain rate (1/s) | 5 | 8.29 | 1.67 | 13 | 8.70 | 1.64 | 19 |
| Radial strain rate (1/s) | 4 | 9.68 | 1.99 | 13 | 9.62 | 2.29 | 19 |

| B. Longitudinal measurements | | Vehicle | | | AngII | | |
|---|---|---|---|---|---|---|---|
| | Segment | Mean | SD | Mean | SD | Mean | SD |
| **Long displacement (mm)** | 1 | 0.554 | 0.123 | 24 | 0.418 | 0.164 | 19 |
| **Long displacement (mm)** | 2 | 0.234 | 0.078 | 24 | 0.195 | 0.102 | 19 |
| **Long displacement (mm)** | 3 | 0.047 | 0.027 | 24 | 0.062 | 0.035 | 19 |
| **Long displacement (mm)** | 6 | 0.033 | 0.023 | 23 | 0.023 | 0.013 | 19 |
| **Long displacement (mm)** | 5 | 0.211 | 0.068 | 24 | 0.144 | 0.093 | 19 |
| **Long displacement (mm)** | 4 | 0.521 | 0.121 | 24 | 0.341 | 0.169 | 19 |
| **Long velocity (cm/s)** | 1 | 1.990 | 0.754 | 24 | 1.290 | 0.618 | 19 |
| **Long velocity (cm/s)** | 2 | 0.931 | 0.373 | 24 | 0.629 | 0.182 | 18 |
| **Long velocity (cm/s)** | 3 | 0.370 | 0.213 | 24 | 0.338 | 0.145 | 19 |
| **Long velocity (cm/s)** | 6 | 0.312 | 0.119 | 24 | 0.407 | 0.223 | 19 |
| **Long velocity (cm/s)** | 5 | 0.933 | 0.440 | 24 | 0.599 | 0.160 | 18 |
| **Long velocity (cm/s)** | 4 | 2.078 | 0.855 | 24 | 1.129 | 0.369 | 18 |
| **Long strain (%)** | 1 | -21.15 | 6.04 | 24 | -17.52 | 7.97 | 19 |
| **Long strain (%)** | 2 | -21.57 | 5.12 | 23 | -19.43 | 6.48 | 19 |
| **Long strain (%)** | 3 | -21.82 | 5.56 | 24 | -23.17 | 6.81 | 19 |
| **Long strain (%)** | 6 | -17.31 | 7.95 | 24 | -16.88 | 6.05 | 19 |
| **Long strain (%)** | 5 | -18.19 | 3.39 | 24 | -15.58 | 4.86 | 19 |
| **Long strain (%)** | 4 | -21.39 | 5.54 | 24 | -14.95 | 6.33 | 19 |
| **Long strain rate (1/s)** | 1 | -8.37 | 3.60 | 24 | -6.14 | 2.81 | 19 |
| **Long strain rate (1/s)** | 2 | -8.12 | 2.60 | 24 | -6.21 | 2.87 | 19 |
| **Long strain rate (1/s)** | 3 | -8.07 | 2.64 | 24 | -7.82 | 2.35 | 19 |
| **Long strain rate (1/s)** | 6 | -6.00 | 2.66 | 24 | -5.58 | 1.50 | 18 |
| **Long strain rate (1/s)** | 5 | -6.68 | 2.19 | 24 | -4.69 | 1.59 | 19 |
| **Long strain rate (1/s)** | 4 | -8.73 | 3.07 | 24 | -5.65 | 2.54 | 19 |
| | | Dabrafenib | | | Dabrafenib/AngII | | |
| | Segment | Mean | SD | Mean | SD | Mean | SD |
| **Long displacement (mm)** | 1 | 0.429 | 0.125 | 13 | 0.444 | 0.174 | 19 |
| **Long displacement (mm)** | 2 | 0.199 | 0.083 | 13 | 0.212 | 0.101 | 19 |
| **Long displacement (mm)** | 3 | 0.067 | 0.052 | 13 | 0.056 | 0.037 | 19 |
| **Long displacement (mm)** | 6 | 0.034 | 0.029 | 13 | 0.025 | 0.014 | 18 |
| **Long displacement (mm)** | 5 | 0.166 | 0.094 | 13 | 0.148 | 0.067 | 19 |
| **Long displacement (mm)** | 4 | 0.404 | 0.146 | 13 | 0.381 | 0.159 | 19 |
| **Long velocity (cm/s)** | 1 | 1.236 | 0.346 | 12 | 1.568 | 0.654 | 19 |
| **Long velocity (cm/s)** | 2 | 0.659 | 0.208 | 13 | 0.770 | 0.318 | 19 |
| **Long velocity (cm/s)** | 3 | 0.386 | 0.199 | 13 | 0.334 | 0.120 | 19 |
| **Long velocity (cm/s)** | 6 | 0.280 | 0.162 | 13 | 0.348 | 0.228 | 19 |
| **Long velocity (cm/s)** | 5 | 0.699 | 0.361 | 13 | 0.732 | 0.264 | 19 |
| **Long velocity (cm/s)** | 4 | 1.384 | 0.636 | 13 | 1.549 | 0.621 | 19 |
| **Long strain (%)** | 1 | -17.70 | 7.45 | 13 | -17.49 | 5.86 | 19 |
| **Long strain (%)** | 2 | -20.10 | 7.13 | 13 | -17.85 | 6.72 | 19 |
| **Long strain (%)** | 3 | -21.18 | 7.95 | 13 | -22.93 | 7.63 | 19 |
| **Long strain (%)** | 6 | -15.92 | 3.02 | 13 | -15.13 | 5.50 | 19 |
| **Long strain (%)** | 5 | -16.37 | 4.65 | 13 | -15.50 | 5.81 | 19 |
| **Long strain (%)** | 4 | -18.17 | 6.03 | 13 | -18.03 | 6.37 | 19 |
| **Long strain rate (1/s)** | 1 | -6.14 | 3.25 | 12 | -7.33 | 3.41 | 19 |
| **Long strain rate (1/s)** | 2 | -5.63 | 1.77 | 11 | -6.46 | 2.35 | 19 |
| **Long strain rate (1/s)** | 3 | -7.33 | 2.67 | 12 | -9.10 | 3.04 | 19 |
| **Long strain rate (1/s)** | 6 | -5.17 | 1.39 | 12 | -5.63 | 2.41 | 19 |
| **Long strain rate (1/s)** | 5 | -4.93 | 1.53 | 11 | -5.25 | 2.30 | 19 |
| **Long strain rate (1/s)** | 4 | -6.25 | 2.63 | 12 | -7.09 | 3.21 | 19 |

**Supplementary Table S7. Global strain measurements for effects of trametinib alone or with dabrafenib (combination) on C57Bl/6J mouse hearts.** Analysis used long axis views of the left ventricle (LV) of echocardiograms collected at 7 d. EDV, end diastolic volume; ESV, end systolic volume; SV, stroke volume; FS, fractional shortening; CO, cardiac output; GLS, global longitudinal strain; EF, ejection fraction; LVEDM, LV end diastolic mass.

| | **Vehicle** | | | **AngII** | | |
|---|---|---|---|---|---|---|
| | **Mean** | **SD** | **N** | **Mean** | **SD** | **N** |
| **EDV (µl)** | 54.40 | 13.11 | 19 | 45.19 | 9.46 | 8 |
| **ESV (µl)** | 22.65 | 6.59 | 19 | 18.95 | 6.83 | 8 |
| **SV (µl)** | 31.75 | 7.87 | 19 | 26.24 | 3.27 | 8 |
| **FS (%)** | 30.18 | 5.49 | 19 | 32.73 | 5.58 | 8 |
| **CO (ml/min)** | 15.87 | 4.01 | 19 | 13.87 | 2.47 | 8 |
| **GLS (%)** | -21.05 | 2.92 | 19 | -17.95 | 4.20 | 8 |
| **EF (%)** | 58.34 | 5.97 | 19 | 58.73 | 7.03 | 8 |
| **LVEDM (mg)** | 57.43 | 5.84 | 19 | 73.42 | 5.61 | 8 |
| **Heart rate (bpm)** | 500 | 42 | 19 | 526 | 40 | 8 |
| | **Trametinib** | | | **Trametinib/AngII** | | |
| | **Mean** | **SD** | **N** | **Mean** | **SD** | **N** |
| **EDV (µl)** | 46.10 | 6.02 | 12 | 45.36 | 6.64 | 9 |
| **ESV (µl)** | 18.86 | 3.30 | 12 | 22.51 | 4.82 | 9 |
| **SV (µl)** | 27.25 | 3.88 | 12 | 22.86 | 4.18 | 9 |
| **FS (%)** | 31.04 | 6.52 | 12 | 25.09 | 4.75 | 9 |
| **CO (ml/min)** | 13.76 | 2.04 | 12 | 11.50 | 2.25 | 9 |
| **GLS (%)** | -19.91 | 3.09 | 12 | -16.28 | 3.15 | 9 |
| **EF (%)** | 58.76 | 4.32 | 12 | 50.27 | 7.52 | 9 |
| **LVEDM (mg)** | 57.83 | 4.95 | 12 | 56.93 | 3.79 | 9 |
| **Heart rate (bpm)** | 507 | 60 | 12 | 505 | 58 | 9 |
| | **Combination** | | | **Combination/AngII** | | |
| **EDV (µl)** | **Mean** | **SD** | **N** | **Mean** | **SD** | **N** |
| **ESV (µl)** | 53.47 | 7.30 | 6 | 43.77 | 8.29 | 9 |
| **SV (µl)** | 21.80 | 6.39 | 6 | 20.72 | 5.70 | 9 |
| **FS (%)** | 31.66 | 1.61 | 6 | 23.05 | 3.61 | 9 |
| **CO (ml/min)** | 29.19 | 3.03 | 6 | 27.32 | 5.15 | 9 |
| **GLS (%)** | 15.87 | 1.39 | 6 | 11.55 | 2.31 | 9 |
| **EF (%)** | -22.84 | 4.12 | 6 | -16.79 | 2.69 | 9 |
| **LVEDM (mg)** | 59.58 | 7.28 | 6 | 52.85 | 6.08 | 9 |
| **Heart rate (bpm)** | 56.33 | 1.35 | 6 | 58.49 | 3.30 | 9 |

**Supplementary Table S8. Segmental strain measurements for effects of trametinib alone or with dabrafenib (combination) on C57Bl/6J mouse hearts.** Analysis used long axis views of the left ventricle (LV). **A**, C57Bl/6J mice treated without inhibitor. **B**, C57Bl/6J mice treated with trametinib. **C**, C57Bl/6J mice treated with trametinib plus dabrafenib (combination). Long, longitudinal.

| **A. No inhibitor** | | **Vehicle** | | | **AngII** | | |
|---|---|---|---|---|---|---|---|
| | **Segment** | **Mean** | **SD** | **Mean** | **SD** | **Mean** | **SD** |
| **Radial displacement (mm)** | 1 | 0.612 | 0.163 | 19 | 0.491 | 0.060 | 8 |
| **Radial displacement (mm)** | 2 | 0.676 | 0.140 | 19 | 0.714 | 0.102 | 8 |
| **Radial displacement (mm)** | 3 | 0.422 | 0.119 | 19 | 0.467 | 0.137 | 8 |
| **Radial displacement (mm)** | 6 | 0.422 | 0.119 | 19 | 0.467 | 0.137 | 8 |
| **Radial displacement (mm)** | 5 | 0.304 | 0.114 | 19 | 0.334 | 0.106 | 8 |
| **Radial displacement (mm)** | 4 | 0.542 | 0.146 | 19 | 0.529 | 0.100 | 8 |
| **Radial velocity (cm/s)** | 1 | 2.019 | 0.460 | 19 | 1.756 | 0.206 | 8 |
| **Radial velocity (cm/s)** | 2 | 2.111 | 0.379 | 19 | 2.065 | 0.222 | 8 |
| **Radial velocity (cm/s)** | 3 | 1.478 | 0.402 | 19 | 1.460 | 0.330 | 8 |
| **Radial velocity (cm/s)** | 6 | 0.669 | 0.313 | 19 | 0.662 | 0.379 | 8 |
| **Radial velocity (cm/s)** | 5 | 0.952 | 0.355 | 19 | 0.937 | 0.308 | 8 |
| **Radial velocity (cm/s)** | 4 | 1.539 | 0.338 | 19 | 1.460 | 0.194 | 8 |
| **Radial strain (%)** | 1 | 39.06 | 12.27 | 19 | 29.908 | 6.868 | 8 |
| **Radial strain (%)** | 2 | 47.52 | 11.61 | 19 | 45.246 | 8.960 | 8 |
| **Radial strain (%)** | 3 | 33.36 | 13.68 | 19 | 32.791 | 8.999 | 8 |
| **Radial strain (%)** | 6 | 28.34 | 11.39 | 19 | 24.584 | 10.489 | 8 |
| **Radial strain (%)** | 5 | 41.54 | 7.78 | 19 | 36.928 | 7.241 | 10 |
| **Radial strain (%)** | 4 | 46.37 | 9.09 | 19 | 38.470 | 8.216 | 8 |
| **Radial strain rate (1/s)** | 1 | 10.07 | 2.14 | 19 | 8.463 | 0.927 | 8 |
| **Radial strain rate (1/s)** | 2 | 12.03 | 2.47 | 19 | 11.390 | 0.980 | 9 |
| **Radial strain rate (1/s)** | 3 | 9.86 | 3.86 | 19 | 8.632 | 1.241 | 8 |
| **Radial strain rate (1/s)** | 6 | 7.69 | 2.31 | 19 | 6.645 | 2.519 | 8 |
| **Radial strain rate (1/s)** | 5 | 10.53 | 2.05 | 19 | 8.938 | 1.384 | 8 |
| **Radial strain rate (1/s)** | 4 | 12.18 | 1.90 | 19 | 10.523 | 2.198 | 8 |
| **Long displacement (mm)** | 1 | 0.598 | 0.135 | 19 | 0.366 | 0.069 | 8 |
| **Long displacement (mm)** | 2 | 0.242 | 0.079 | 19 | 0.153 | 0.038 | 8 |
| **Long displacement (mm)** | 3 | 0.037 | 0.019 | 18 | 0.055 | 0.038 | 8 |
| **Long displacement (mm)** | 6 | 0.041 | 0.038 | 19 | 0.027 | 0.016 | 8 |
| **Long displacement (mm)** | 5 | 0.254 | 0.068 | 19 | 0.138 | 0.075 | 8 |
| **Long displacement (mm)** | 4 | 0.556 | 0.141 | 19 | 0.328 | 0.109 | 8 |
| **Long velocity (cm/s)** | 1 | 1.869 | 0.590 | 19 | 1.336 | 0.502 | 8 |
| **Long velocity (cm/s)** | 2 | 0.833 | 0.262 | 19 | 0.594 | 0.106 | 8 |
| **Long velocity (cm/s)** | 3 | 0.430 | 0.256 | 19 | 0.483 | 0.172 | 8 |
| **Long velocity (cm/s)** | 6 | 0.277 | 0.111 | 19 | 0.312 | 0.165 | 8 |
| **Long velocity (cm/s)** | 5 | 0.935 | 0.295 | 19 | 0.705 | 0.322 | 8 |
| **Long velocity (cm/s)** | 4 | 1.942 | 0.661 | 19 | 1.418 | 0.659 | 8 |
| **Long strain (%)** | 1 | -22.53 | 6.93 | 19 | -12.61 | 4.88 | 8 |
| **Long strain (%)** | 2 | -24.94 | 6.70 | 19 | -22.21 | 5.26 | 8 |
| **Long strain (%)** | 3 | -22.65 | 5.20 | 19 | -21.20 | 4.72 | 7 |
| **Long strain (%)** | 6 | -15.43 | 4.86 | 19 | -14.97 | 8.49 | 8 |
| **Long strain (%)** | 5 | -18.78 | 3.43 | 19 | -15.25 | 5.12 | 8 |
| **Long strain (%)** | 4 | -19.65 | 5.17 | 19 | -16.36 | 6.59 | 8 |
| **Long strain rate (1/s)** | 1 | -9.12 | 3.71 | 19 | -4.89 | 1.72 | 8 |
| **Long strain rate (1/s)** | 2 | -8.92 | 2.64 | 19 | -7.74 | 2.28 | 8 |
| **Long strain rate (1/s)** | 3 | -8.62 | 2.96 | 19 | -7.79 | 2.85 | 8 |
| **Long strain rate (1/s)** | 6 | -5.61 | 2.26 | 19 | -5.38 | 2.88 | 8 |
| **Long strain rate (1/s)** | 5 | -6.41 | 1.86 | 19 | -5.20 | 1.69 | 8 |
| **Long strain rate (1/s)** | 4 | -7.99 | 2.51 | 19 | -6.95 | 2.93 | 8 |

| B. Plus trametinib | | Vehicle | | | AngII | | |
|---|---|---|---|---|---|---|---|
| | Segment | Mean | SD | Mean | SD | Mean | SD |
| Radial displacement (mm) | 1 | 0.568 | 0.121 | 12 | 0.456 | 0.148 | 9 |
| Radial displacement (mm) | 2 | 0.699 | 0.107 | 12 | 0.537 | 0.134 | 9 |
| Radial displacement (mm) | 3 | 0.405 | 0.128 | 12 | 0.358 | 0.069 | 9 |
| Radial displacement (mm) | 6 | 0.405 | 0.128 | 12 | 0.358 | 0.069 | 9 |
| Radial displacement (mm) | 5 | 0.276 | 0.086 | 12 | 0.206 | 0.084 | 9 |
| Radial displacement (mm) | 4 | 0.542 | 0.068 | 12 | 0.450 | 0.119 | 9 |
| Radial velocity (cm/s) | 1 | 1.871 | 0.355 | 12 | 1.569 | 0.393 | 9 |
| Radial velocity (cm/s) | 2 | 2.150 | 0.338 | 12 | 1.670 | 0.339 | 9 |
| Radial velocity (cm/s) | 3 | 1.571 | 0.403 | 12 | 1.167 | 0.308 | 9 |
| Radial velocity (cm/s) | 6 | 0.762 | 0.329 | 12 | 0.610 | 0.263 | 9 |
| Radial velocity (cm/s) | 5 | 0.897 | 0.206 | 12 | 0.641 | 0.257 | 9 |
| Radial velocity (cm/s) | 4 | 1.578 | 0.255 | 12 | 1.242 | 0.344 | 9 |
| Radial strain (%) | 1 | 33.31 | 9.19 | 12 | 29.592 | 10.685 | 9 |
| Radial strain (%) | 2 | 49.01 | 6.75 | 12 | 38.325 | 11.600 | 9 |
| Radial strain (%) | 3 | 36.94 | 10.51 | 12 | 26.081 | 3.796 | 9 |
| Radial strain (%) | 6 | 28.08 | 11.90 | 12 | 19.819 | 8.776 | 9 |
| Radial strain (%) | 5 | 36.25 | 7.85 | 12 | 27.287 | 9.959 | 9 |
| Radial strain (%) | 4 | 41.95 | 4.77 | 12 | 37.262 | 9.708 | 9 |
| Radial strain rate (1/s) | 1 | 8.45 | 1.89 | 12 | 7.967 | 2.177 | 9 |
| Radial strain rate (1/s) | 2 | 11.78 | 1.69 | 11 | 9.377 | 2.087 | 9 |
| Radial strain rate (1/s) | 3 | 11.25 | 3.29 | 12 | 7.857 | 1.772 | 9 |
| Radial strain rate (1/s) | 6 | 8.02 | 2.31 | 12 | 6.050 | 1.656 | 9 |
| Radial strain rate (1/s) | 5 | 9.59 | 2.05 | 12 | 7.591 | 1.696 | 9 |
| Radial strain rate (1/s) | 4 | 11.26 | 2.09 | 12 | 9.324 | 2.459 | 9 |
| Long displacement (mm) | 1 | 0.492 | 0.154 | 12 | 0.396 | 0.158 | 9 |
| Long displacement (mm) | 2 | 0.228 | 0.045 | 6 | 0.160 | 0.074 | 9 |
| Long displacement (mm) | 3 | 0.054 | 0.034 | 6 | 0.052 | 0.039 | 9 |
| Long displacement (mm) | 6 | 0.070 | 0.023 | 6 | 0.025 | 0.025 | 9 |
| Long displacement (mm) | 5 | 0.265 | 0.062 | 6 | 0.139 | 0.084 | 9 |
| Long displacement (mm) | 4 | 0.510 | 0.165 | 6 | 0.358 | 0.172 | 9 |
| Long velocity (cm/s) | 1 | 1.608 | 0.430 | 6 | 1.451 | 0.602 | 9 |
| Long velocity (cm/s) | 2 | 0.795 | 0.167 | 6 | 0.624 | 0.290 | 9 |
| Long velocity (cm/s) | 3 | 0.358 | 0.127 | 6 | 0.353 | 0.076 | 9 |
| Long velocity (cm/s) | 6 | 0.285 | 0.041 | 6 | 0.283 | 0.166 | 9 |
| Long velocity (cm/s) | 5 | 0.810 | 0.170 | 7 | 0.759 | 0.258 | 9 |
| Long velocity (cm/s) | 4 | 1.679 | 0.384 | 6 | 1.443 | 0.680 | 9 |
| Long strain (%) | 1 | -21.56 | 1.99 | 6 | -14.90 | 4.88 | 9 |
| Long strain (%) | 2 | -23.54 | 1.80 | 6 | -18.22 | 6.71 | 9 |
| Long strain (%) | 3 | -21.40 | 7.56 | 6 | -19.34 | 4.79 | 9 |
| Long strain (%) | 6 | -19.98 | 5.31 | 6 | -13.68 | 6.56 | 9 |
| Long strain (%) | 5 | -15.23 | 5.41 | 6 | -12.53 | 3.76 | 9 |
| Long strain (%) | 4 | -18.60 | 5.93 | 6 | -17.26 | 6.35 | 9 |
| Long strain rate (1/s) | 1 | -7.66 | 0.67 | 6 | -5.64 | 2.56 | 9 |
| Long strain rate (1/s) | 2 | -8.47 | 1.50 | 6 | -6.47 | 2.42 | 9 |
| Long strain rate (1/s) | 3 | -9.55 | 3.84 | 6 | -7.27 | 2.26 | 9 |
| Long strain rate (1/s) | 6 | -7.00 | 2.43 | 6 | -4.35 | 1.74 | 9 |
| Long strain rate (1/s) | 5 | -5.25 | 1.79 | 6 | -4.43 | 2.12 | 9 |
| Long strain rate (1/s) | 4 | -7.03 | 2.07 | 6 | -6.27 | 2.19 | 9 |

| C. Plus combination | | Vehicle | | | AngII | | |
|---|---|---|---|---|---|---|---|
| | Segment | Mean | SD | Mean | SD | Mean | SD |
| Radial displacement (mm) | 1 | 0.586 | 0.072 | 6 | 0.461 | 0.133 | 9 |
| Radial displacement (mm) | 2 | 0.707 | 0.160 | 6 | 0.559 | 0.086 | 9 |
| Radial displacement (mm) | 3 | 0.469 | 0.135 | 6 | 0.358 | 0.116 | 9 |
| Radial displacement (mm) | 6 | 0.469 | 0.135 | 6 | 0.358 | 0.116 | 9 |
| Radial displacement (mm) | 5 | 0.335 | 0.094 | 6 | 0.250 | 0.104 | 9 |
| Radial displacement (mm) | 4 | 0.491 | 0.101 | 6 | 0.434 | 0.122 | 9 |
| Radial velocity (cm/s) | 1 | 1.975 | 0.171 | 6 | 1.635 | 0.334 | 9 |
| Radial velocity (cm/s) | 2 | 2.234 | 0.450 | 6 | 1.836 | 0.205 | 9 |
| Radial velocity (cm/s) | 3 | 1.748 | 0.562 | 6 | 1.285 | 0.290 | 9 |
| Radial velocity (cm/s) | 6 | 0.978 | 0.342 | 6 | 0.528 | 0.202 | 9 |
| Radial velocity (cm/s) | 5 | 1.054 | 0.233 | 6 | 0.719 | 0.261 | 9 |
| Radial velocity (cm/s) | 4 | 1.398 | 0.278 | 6 | 1.272 | 0.268 | 9 |
| Radial strain (%) | 1 | 36.35 | 5.38 | 6 | 29.956 | 10.519 | 9 |
| Radial strain (%) | 2 | 50.10 | 8.21 | 6 | 39.877 | 7.326 | 9 |
| Radial strain (%) | 3 | 42.56 | 13.75 | 6 | 24.133 | 10.295 | 9 |
| Radial strain (%) | 6 | 36.45 | 14.26 | 6 | 20.684 | 11.034 | 9 |
| Radial strain (%) | 5 | 41.46 | 11.19 | 6 | 33.608 | 9.134 | 9 |
| Radial strain (%) | 4 | 38.30 | 8.30 | 6 | 34.943 | 8.992 | 9 |
| Radial strain rate (1/s) | 1 | 9.27 | 1.69 | 6 | 7.930 | 2.155 | 9 |
| Radial strain rate (1/s) | 2 | 12.21 | 2.30 | 6 | 9.839 | 1.283 | 9 |
| Radial strain rate (1/s) | 3 | 10.99 | 2.93 | 6 | 7.795 | 1.860 | 9 |
| Radial strain rate (1/s) | 6 | 10.52 | 3.76 | 6 | 6.044 | 2.257 | 9 |
| Radial strain rate (1/s) | 5 | 10.36 | 1.69 | 6 | 8.258 | 1.834 | 9 |
| Radial strain rate (1/s) | 4 | 9.19 | 2.06 | 6 | 9.875 | 2.829 | 9 |
| Long displacement (mm) | 1 | 0.548 | 0.111 | 6 | 0.418 | 0.099 | 9 |
| Long displacement (mm) | 2 | 0.184 | 0.062 | 6 | 0.175 | 0.047 | 9 |
| Long displacement (mm) | 3 | 0.060 | 0.030 | 6 | 0.045 | 0.034 | 9 |
| Long displacement (mm) | 6 | 0.035 | 0.019 | 6 | 0.012 | 0.011 | 9 |
| Long displacement (mm) | 5 | 0.174 | 0.073 | 6 | 0.129 | 0.079 | 9 |
| Long displacement (mm) | 4 | 0.381 | 0.149 | 6 | 0.352 | 0.109 | 9 |
| Long velocity (cm/s) | 1 | 1.329 | 0.333 | 6 | 1.322 | 0.391 | 9 |
| Long velocity (cm/s) | 2 | 0.757 | 0.256 | 6 | 0.619 | 0.202 | 9 |
| Long velocity (cm/s) | 3 | 0.340 | 0.068 | 6 | 0.283 | 0.120 | 9 |
| Long velocity (cm/s) | 6 | 0.189 | 0.070 | 5 | 0.205 | 0.116 | 9 |
| Long velocity (cm/s) | 5 | 0.671 | 0.297 | 5 | 0.607 | 0.284 | 9 |
| Long velocity (cm/s) | 4 | 1.447 | 0.374 | 6 | 1.360 | 0.569 | 9 |
| Long strain (%) | 1 | -20.89 | 4.13 | 6 | -15.34 | 6.53 | 9 |
| Long strain (%) | 2 | -22.08 | 3.94 | 6 | -19.48 | 6.69 | 9 |
| Long strain (%) | 3 | -26.58 | 8.12 | 6 | -20.60 | 8.98 | 9 |
| Long strain (%) | 6 | -16.47 | 6.85 | 6 | -13.00 | 7.48 | 9 |
| Long strain (%) | 5 | -13.08 | 3.81 | 6 | -13.19 | 3.20 | 9 |
| Long strain (%) | 4 | -19.39 | 6.66 | 6 | -16.91 | 4.33 | 9 |
| Long strain rate (1/s) | 1 | -8.27 | 1.45 | 6 | -6.11 | 2.51 | 9 |
| Long strain rate (1/s) | 2 | -6.64 | 1.76 | 6 | -6.78 | 2.45 | 9 |
| Long strain rate (1/s) | 3 | -10.09 | 2.46 | 6 | -7.64 | 2.92 | 9 |
| Long strain rate (1/s) | 6 | -5.46 | 1.73 | 6 | -4.32 | 2.28 | 9 |
| Long strain rate (1/s) | 5 | -4.63 | 1.12 | 6 | -4.69 | 1.55 | 9 |
| Long strain rate (1/s) | 4 | -6.96 | 1.68 | 6 | -6.34 | 1.81 | 9 |

**Supplementary Figure S1. Segmental strain data for effects of dabrafenib on AngII-induced hypertrophy: scatter plots for peak values.** Male C57Bl/6J mice (10 wks) were treated with vehicle only (Veh), 3 mg/kg/d dabrafenib (Dab), 0.8 mg/kg/d AngII or AngII with dabrafenib, and echocardiograms were collected at 7 d. Long axis B-mode images were analysed using speckle-tracking; data for individual segments were exported for analysis. Data are for segmental strain for peak radial (**A**) or longitudinal (**B**) endocardial displacement, velocity, strain and strain rate. Individual datapoints are shown with the means ± 95% CI. Statistical analysis used 2-way ANOVA with Tukey's post-test. Ssignificant differences (p<0.05) are in bold type.

**A**

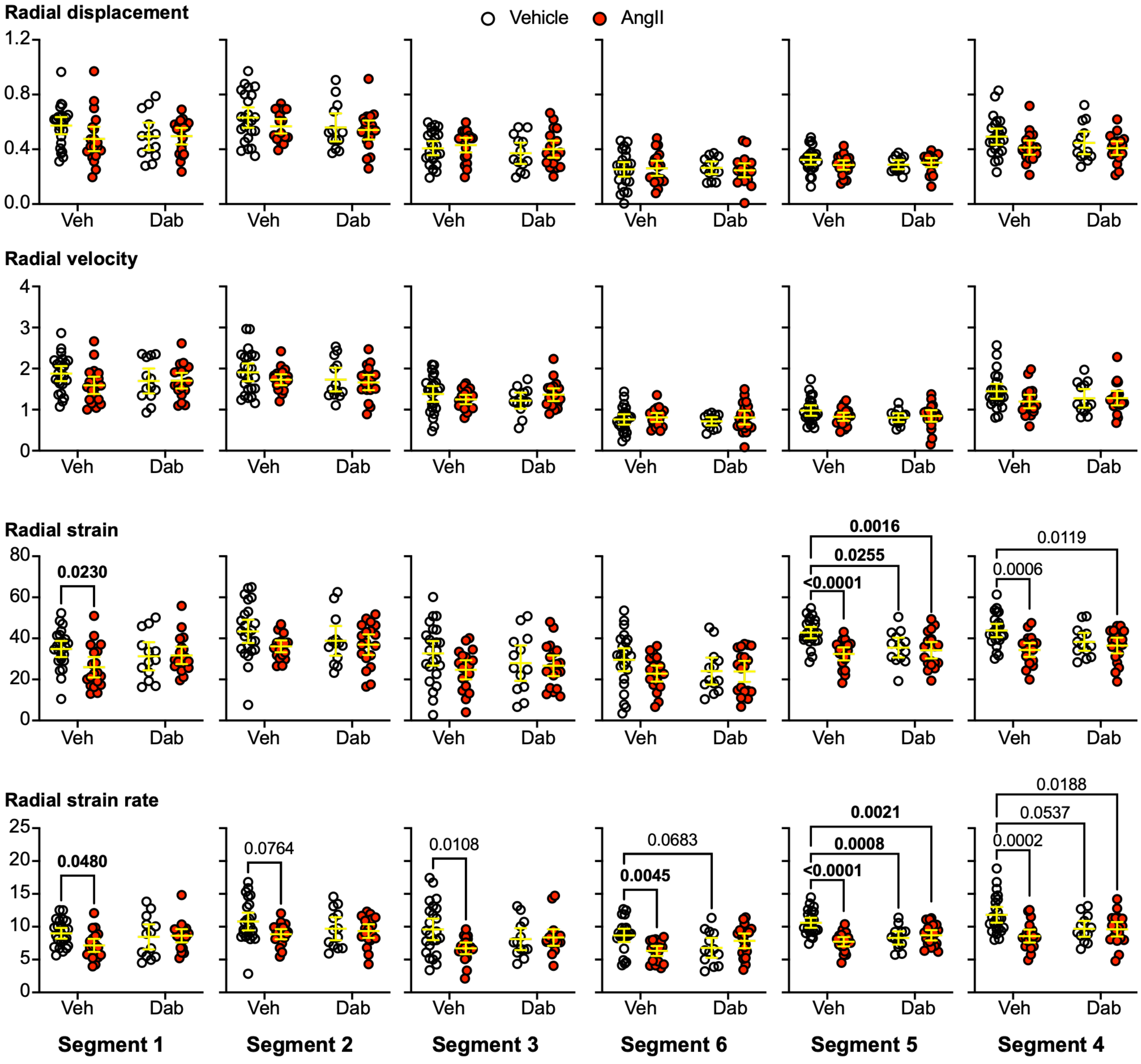

**B**

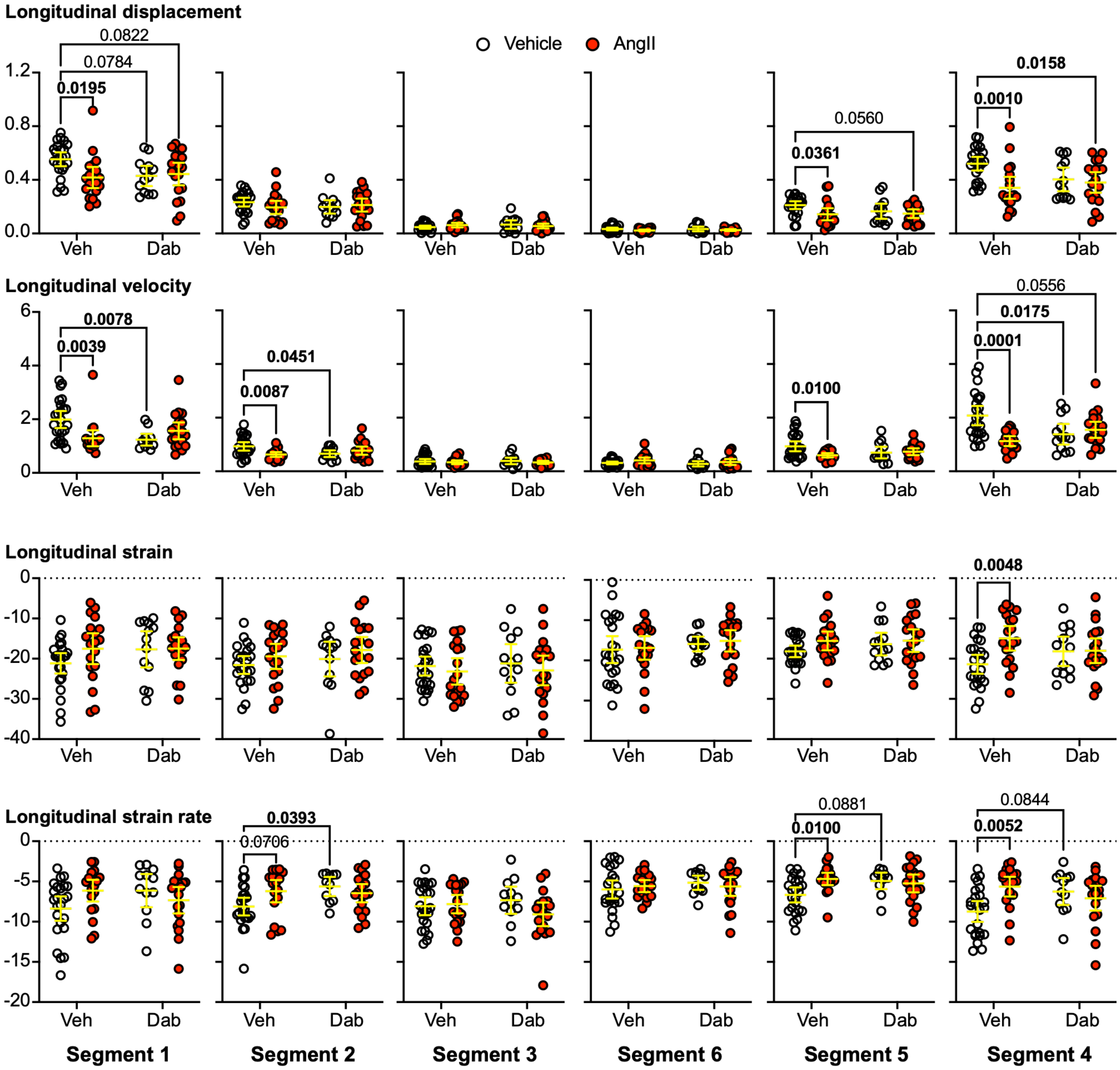
Longitudinal displacement
Vehicle
AngII
0.0822
0.0784
0.0195
0.0560
0.0361
0.0158
0.0010
Longitudinal velocity
0.0078
0.0039
0.0451
0.0087
0.0100
0.0556
0.0175
0.0001
Longitudinal strain
0.0048
Longitudinal strain rate
0.0393
0.0706
0.0881
0.0100
0.0844
0.0052
Veh
Dab
Segment 1
Segment 2
Segment 3
Segment 6
Segment 5
Segment 4

**Supplementary Figure S2. Segmental strain data for effects of trametinib or trametinib with dabrafenib on AngII-induced hypertrophy: scatter plots for peak values.** Male C57Bl/6J mice (10 wks) were treated with vehicle (Veh), 1 mg /kg/d trametinib alone (T) or with 3 mg/kg/d dabrafenib (C) in the absence or presence of 0.8 mg/kg/d AngII, and echocardiograms were collected at 7 d. Long axis B-mode images were analysed using speckle-tracking;data for individual segments were exported for analysis. Data are for segmental strain for peak radial (**A**) or longitudinal (**B**) endocardial displacement, velocity, strain and strain rate. Individual datapoints are shown with the means ± 95% CI. Statistical analysis used 2-way ANOVA with Tukey's post-test. Significant differences (p<0.05) are in bold type.

**A**

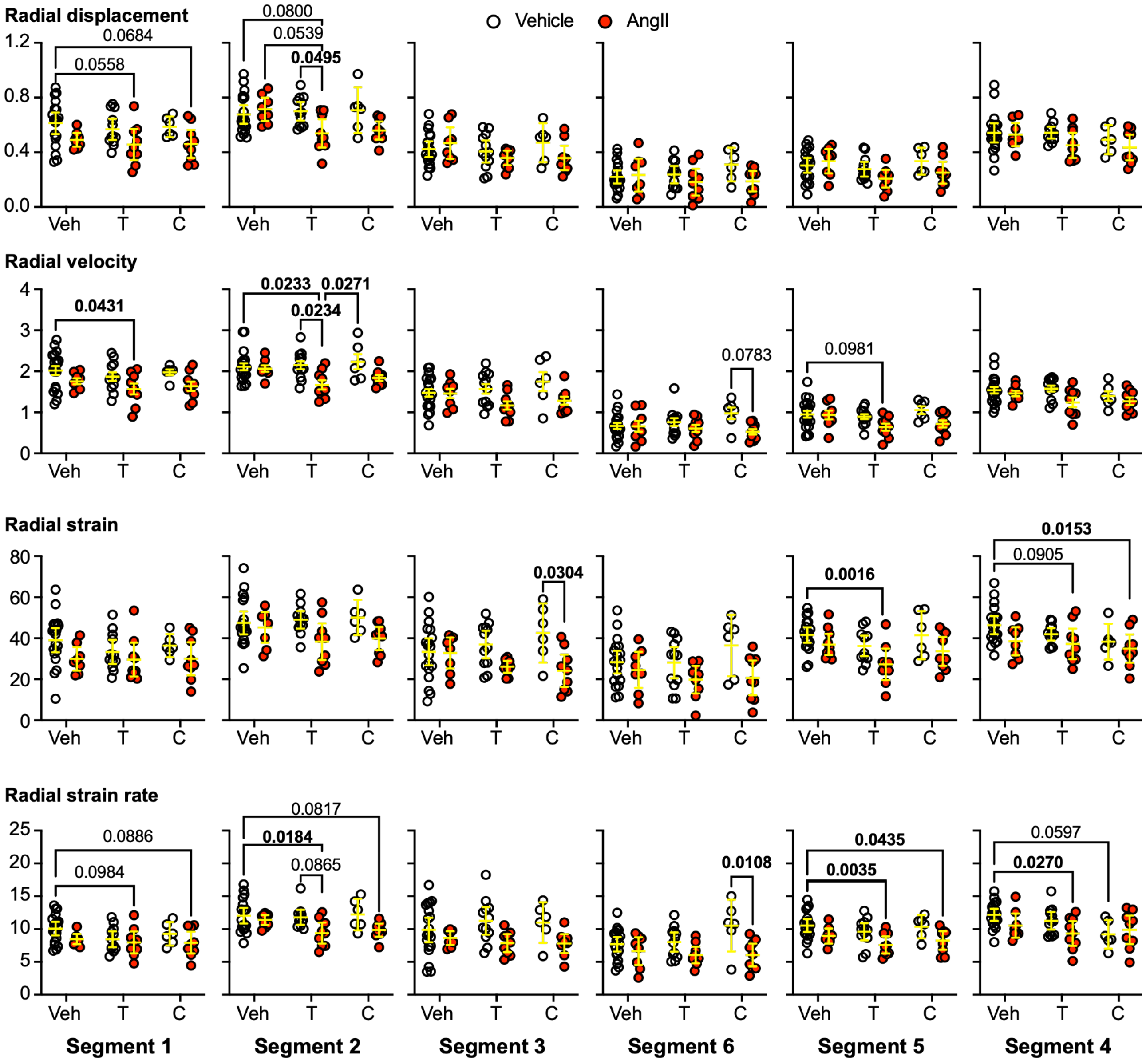

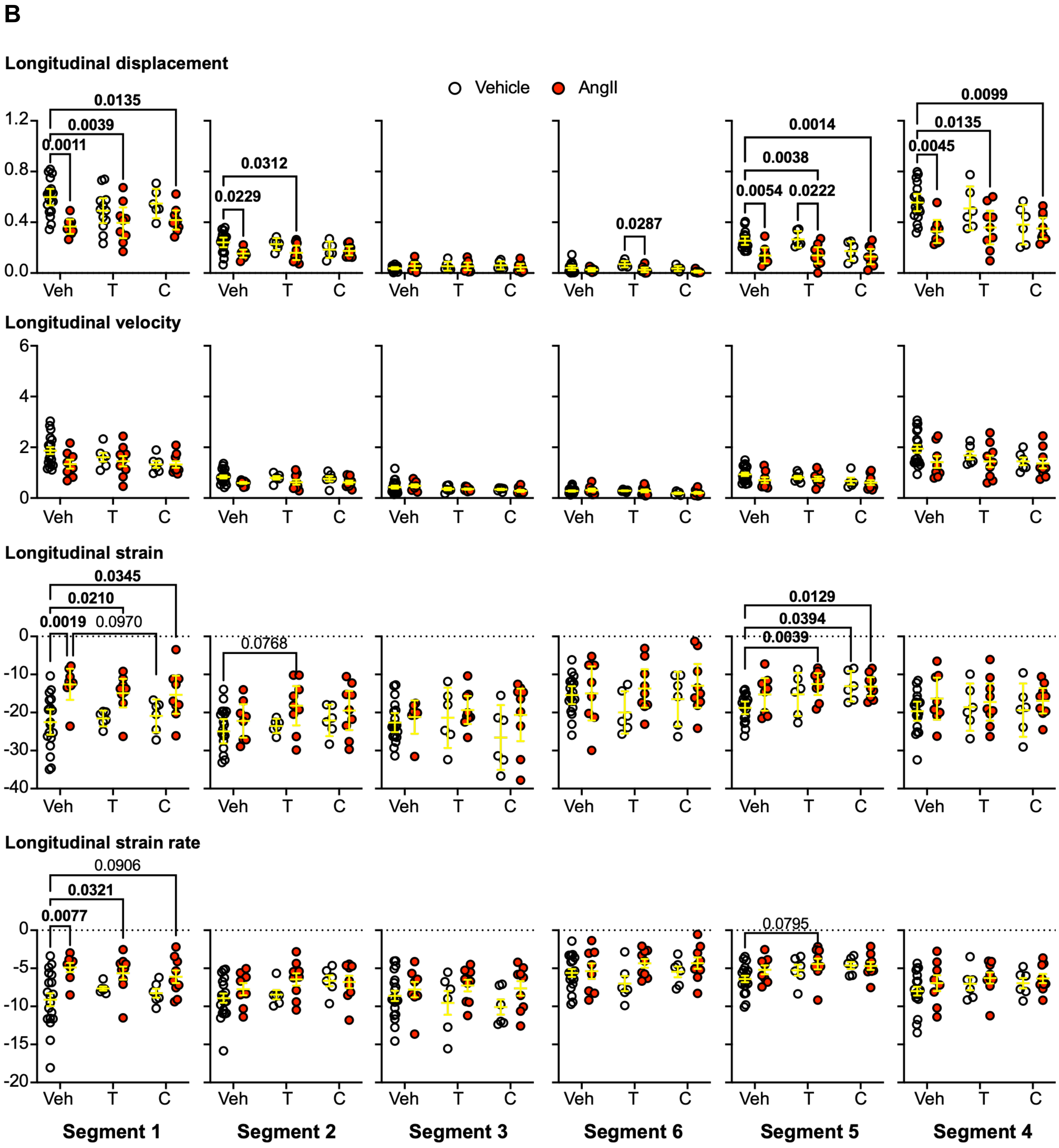
B
Longitudinal displacement
Vehicle
AngII
Longitudinal velocity
Longitudinal strain
Longitudinal strain rate
Veh
T
C
Segment 1
Segment 2
Segment 3
Segment 6
Segment 5
Segment 4
0.0135
0.0039
0.0011
0.0312
0.0229
0.0287
0.0014
0.0038
0.0054
0.0222
0.0099
0.0135
0.0045
0.0345
0.0210
0.0019
0.0970
0.0768
0.0129
0.0394
0.0039
0.0906
0.0321
0.0077
0.0795
